# Charting the Cannabis plant chemical space with computational metabolomics

**DOI:** 10.1101/2024.02.21.581363

**Authors:** Akhona Myoli, Mpho Choene, Abidemi Paul Kappo, Ntakadzeni Edwin Madala, Justin J.J. van der Hooft, Fidele Tugizimana

## Abstract

**Introduction:** The chemical classification of Cannabis is typically confined to the cannabinoid content, whilst Cannabis encompasses diverse chemical classes that vary in abundance among all its varieties. Hence, neglecting other chemical classes within Cannabis strains results in a restricted and biased comprehension of elements that may contribute to chemical intricacy and the resultant medicinal qualities of the plant.

**Objectives:** Thus, herein, we report a computational metabolomics study to elucidate the Cannabis metabolic map beyond the cannabinoids.

**Methods:** Mass spectrometry-based computational tools were used to mine and evaluate the methanolic leaf and flower extracts of two Cannabis cultivars: Amnesia haze (AMNH) and Royal dutch cheese (RDC).

**Results:** The results revealed the presence of different chemical compound classes including cannabinoids, but extending it to flavonoids, polyketides, and phospholipids at varying distributions across the cultivar plant tissues. Therefore, the two cultivars were differentiated based on the overall chemical content of their plant tissues where AMNH was observed to be more dominant in the flavonoid content while RDC was more dominant in the lipid-like molecules. Additionally, *in silico* molecular docking studies in combination with biological assay studies indicated the potentially differing anti-cancer properties of the two cultivars resulting from the elucidated chemical profiles.

**Conclusion:** These findings highlight distinctive chemical profiles beyond cannabinoids in Cannabis strains. This novel mapping of the metabolomic landscape of Cannabis provides actionable insights into plant biochemistry and justifies selecting certain varieties for medicinal use.

## 1. Introduction

The Cannabis plant is of the Cannabaceae family and is known to grow in various forms of cultivars or strains. Cannabis is popularly known for both its recreational use and there is emerging awareness of its medicinal properties (Cicaloni et al., 2022). There are currently more than 700 strains of Cannabis with trivial names such as Purple Kush, White Widow, Amnesia Haze and Royal Dutch cheese. When drawing focus on the chemical composition of the numerous Cannabis strains or cultivars, the chemical classification or differentiation is limited to several categories or chemovars which are based only on the plant’s cannabinoid content (Gloss, 2015; Lewis et al., 2018). The cannabinoid chemical class comprises a group of compounds that naturally occur in Cannabis varieties. The chemical structure of these compounds is characterized by a C21 terpene phenolic backbone, which can be traced in parent cannabinoids, cannabinoid derivatives, and transformation products. These cannabinoids are further divided into sub-classes, namely; cannabichromene (CBC), cannabidiol (CBD), cannabielsoin (CBE), cannabigerol (CBG), cannabicyclol (CBL), cannabinol (CBN), cannabinodiol (CBND), cannabitriol (CBT), △^8^-trans-tetrahydrocannabinol (△^8^-THC), △^9^-trans-tetrahydrocannabinol (△^9^-THC), and miscellaneous-type cannabinoids (Filipiuc et al., 2021; Radwan et al., 2021; Procaccia et al., 2022).

Type I Cannabis cultivars are characterized by high tetrahydrocannabinol (THC) levels and are used for both medicinal and recreational purposes. Type III Cannabis contains high levels of cannabidiol (CBD), which has been acknowledged for its therapeutic benefits more than type I. This has also led to the emergence of additional chemovars such as type II Cannabis which is known to have equal levels of both THC and CBD compounds (Lewis et al., 2018). Nonetheless, some of the reported medicinal applications and benefits of Cannabis include alleviating nausea and pain in cancer patients receiving chemotherapy, and alleviating neurological symptoms such as anxiety, stress and sleeping problem (Moltke & Hindocha, 2021). Additionally, studies have shown promising evidence of the therapeutic effects of the Cannabis plant (including drugs and essential oils derived from the plant) in the suppression of diseases such as cancer, Alzheimer’s disease, and Huntington’s disease (Kogan & Mechoulam, 2007; Odieka et al., 2022). Although Cannabis is expected to have a rich, complex, and diverse chemical composition, however, so far, the only bioactive chemical constituents that have been well-studied from Cannabis are cannabinoids (Sarma et al., 2020).

The cannabinoid chemical class is distributed in varying amounts across all the different plant parts of Cannabis. The typical anatomy of the Cannabis plant is comprised of the leaves, flowers, stem, and roots. When considering the plant parts or plant tissues of Cannabis, cannabinoids are widely known to be abundant in the flowers (inflorescence) of the plant compared to the other plant tissues (Jin et al., 2020). Consequently, several pharmaceutical companies (supported by numerous scientific studies) exclusively focus on isolated compounds or crude extracts from Cannabis flowers (inflorescence) for the development of new drugs due to the high cannabinoid content found in the flowers(Sarma et al., 2020; Balant et al., 2021;). However, research shows that the use of leaf or seed in traditional medicine (Cannabis) is often more important than the use of inflorescence for the treatment of certain ailments (Balant et al., 2021). This suggests that Cannabis or its other plant tissues can be alternative sources for drug discovery. Therefore, the comprehensive exploration of the full phytochemical space of Cannabis and its different plant parts remains important as this can aid in identifying Cannabis cultivars or specific plant parts that are more useful for specific ailments (Aizpurua-Olaizola et al., 2016; Namdar et al., 2018; Balant et al., 2021).

Nevertheless, the cannabinoid-centred approach continues to dominate the research field, and this has limited the exploration of the full potential of Cannabis chemistry. Furthermore, with an increasing attention to Cannabis, globally, and a growing momentum for legalization and commercialization of the plant (in some countries this is currently ongoing), there is a need to comprehensively characterize the chemistry of Cannabis (and its various plant tissues) beyond cannabinoids to understand its full chemical composition (Lowe et al., 2021; Pattnaik et al., 2022). The increasing scientific efforts to characterize and explore the metabolome of Cannabis encompass the utilization of omics sciences, particularly metabolomics, which provides distinctive avenues to investigate the plant’s metabolism and unravel the biochemical mechanisms driving the synthesis of various specialized metabolites within its tissues. Furthermore, metabolomics can facilitate and accelerate the search for novel bioactive compounds from plant crude extracts (Aliferis & Bernard-Perron, 2020; Vásquez-Ocmín et al., 2021; C. R. Li et al., 2022).

The application of metabolomics in Cannabis research and development (R&D), coined the term “cannabolomics”, is still in its infancy and there is a need for the optimization of bioanalytical protocols, instrument analysis and metabolite databases to improve the robustness of this approach (Aliferis & Bernard-Perron, 2020). Nonetheless, cannabolomics has been suggested for the classification of Cannabis into chemical varieties or “chemovars,” to emphasize the unique overall biochemical profile of the Cannabis plant of interest (Ladha et al., 2020). Such chemical profiling or metabolic phenotyping can confirm the composition and quality of the chemovar of interest, with variations indicating its medicinal applications. Thus, the study reported herein is a computational metabolomics work to elucidate a Cannabis metabolomic atlas of two cultivars for which sufficient seed material was available: Amnesia Haze and Royal Dutch Cheese cultivars (both type I chemovars). We mapped their metabolome content beyond cannabinoids using a suite of computational metabolomics strategies including feature-based molecular networking, substructural discovery method (MS2LDA), *in silico* tools (*e.g*., NAP and DEREPLICTOR+), and MolNetEnhancer. The study contributes to ongoing Cannabis R&D efforts, particularly the comprehensive characterization of the plant chemistries of these two drug-type cultivars (a first of its kind) that have not been investigated or discussed in literature but form part of the many acclaimed Cannabis strains that have been generally hailed for their therapeutic properties against diseases such cancer. Hence, this study is also part of the ongoing efforts to determine the anti-cancer properties of Cannabis chemovars and aims to contribute and build on the evaluation of the bioactivity elicited by the varying chemical composition of Cannabis chemovars.

## 2. Experimental procedure

### 2.1 Plant cultivation and harvesting

Cannabis seeds, Amnesia haze (a hybrid genetically comprised of 80% *C. sativa* & 20% *C. indica*) and Royal dutch cheese (a hybrid genetically comprised of 70% *C. indica* & 30% *C. sativa*), were purchased from Marijuana SA (Pty) Ltd (Cape town, South Africa). The seeds were grown into plants in Freedom farms premium classic growing medium (Freedom Farms Horticulture Technologies, Cape town, South Africa) at 24 °C, 70% humidity and a 24-hour light cycle for germination and vegetative stages. The flowering stage for both cultivars was initiated at 27 °C, 30% humidity and a 12-hour light/12-hour dark cycle. Seven-week-old plants in their vegetative stages were harvested for leaves per cultivar and 16-week-old plants in their flowering stages were harvested for flowers. The harvested plant-tissues were freeze-dried and crushed with a blender to powder form and stored at room temperature until metabolite extraction.

### 2.2 Metabolite extraction and standard preparation

Fifty milligrams (50 mg) of the samples were weighed and extracted in 1 mL of 80% methanol (Chemlab, UK). The samples (leaves and flowers per respective cultivar) were spun overnight in a digital rotisserie tube rotator at 70 rpm. The crude extracts were then centrifuged at 158 × *g* in a benchtop fixed-angle centrifuge (Thermo Fisher, Johannesburg, South Africa). After centrifugation, the supernatants were filtered using 0.22 µm nylon filters into glass vials with 500 µL inserts. In this study, seven independent replicates for each sample group were weighed and prepared. The prepared samples were then stored at 4 °C until analysis. For chemical baiting, the vitexin standard was prepared in 50% methanol to a concentration of 5 ppm. The standard was treated in the same manner as the samples for experimental procedures.

### 2.3 Data acquisition – LC-MS/MS analyses of crude extracts

Samples (methanol extracts) were analyzed on a liquid chromatography–quadrupole time-of-flight mass spectrometry (LC-qToF-MS) instrument (LCMS-9030, Shimadzu Corporation, Kyoto, Japan). For chromatographic separation, a sample volume of 3 µL was injected on a Shim-pack C18 column (100 mm × 2.1 mm, 2.7 µm) (Shimadzu Corporation, Kyoto, Japan) thermostatted at 55 °C. In addition to the stationary reverse phase, the chromatography was carried out with a binary mobile phase, applying a gradient elution method. The solvent system comprised solvent A consisting of 0.1% formic acid in Milli-Q water (both HPLC grade, Merck, Darmstadt, Germany) and solvent B being methanol (UHPLC grade, Romil SpS, Cambridge, UK) with 0.1% formic acid, with a flow rate of 0.4 mL/min. The gradient elution was performed as follows, B referring to organic composition (*i.e*., solvent B): 5% B for 3 min, 5–40% B over 3–5 min, 40-95% B over 5-12 min, then 95% B for from 12 min to 18 min. The gradient was changed back to initial 5% B at 18 min and kept to 20 min. The column was re-equilibrated for 3 min.

The effluents from chromatographic separation were further analyszd with the high-definition mass spectrometer, equipped with electrospray ionization (ESI), acquiring both negative and positive spectral data. The mass spectrometer parameters used were the following: 4.5 kV interface voltage, with interface temperature of 300 °C; 3 L/min flow rate for nebulization and dry gas; DL temperature of 250 °C, and 400 °C for heat block; detector was operated at 1.8 kV voltage. For monitoring the accuracy of acquired mass-to-charge ratio (*m/z*), sodium iodide (NaI) was used as a calibration solution. Both non-fragmented (MS1) and fragmented (MS2) spectral data were acquired with *m/z* range of 100–1200 Da. For MS/MS experiments, data-dependent acquisition method was applied, with an intensity threshold of 5000 counts. The collision-induced dissociation (CID) method was applied for the fragmentation of ions, using argon as a collision gas at a collision energy of 30 eV with a spread of 5 eV.

### 2.4 Data mining: processing and chemometrics analyses

The acquired raw datasets (ESI negative and positive centroid data) were processed using MetaboAnalyst version 5.0 (Pang et al., 2021). The data was processed with UPLC-Q/TOF default parameters using the centWave method. From the default setting, the max peak width was changed to 15. The resulting feature tables with 3560 and 3570 features for (ESI) negative and positive datasets respectively were imported into soft independent modelling of class analogy (SIMCA) version 17.0 software (Sartorius, South Africa) for chemometrics modelling. Principal component analysis (PCA) and hierarchical cluster analysis (HCA) models were computed for data exploration, and group classification and discriminant analysis.

### 2.5 Molecular networking and Metabolite Annotation

Prior to computing the feature-based molecular networks (FBMN) through the Global Natural Product Social (GNPS), the acquired raw datasets obtained from the Shimadzu LCMS-9030-qTOF-MS were converted to open-source format (.mzML) files. The mzML files were then uploaded to Mass Spectrometry-Data Independent AnaLysis (MS-DIAL) platform for data processing. The parameters used for MS-DIAL data processing parameters included mass accuracy MS1 and MS2 tolerance of 0.25 Da and 0.1 Da respectively, with the MS/MS range of 50-800 Da; the minimum peak height was 2000 amplitude, mass slice width of 0.1 Da for peak detection, a sigma window value of 0.5 was used, and retention time tolerance was 0.1 min and MS1 tolerance set at 0.015.

Post MS-DIAL data processing, the GNPS export files, both GNPS MGF files and feature quantification tables, were exported into the GNPS ecosystem using the WinSCP server for molecular networking. FBMNs were then computed and generated using the GNPS molecular networking workflow. The parameters used for computing FBMN for the leaves and flower spectral datasets included precursor ion mass tolerance of 0.25 Da with fragment ion mass tolerance of 0.25 Da. For spectral similarity, a cosine score cut off was 0.65, with a minimum of 4 matched fragment ions. For annotation, various spectral libraries were searched and these included MassBank, ReSpect and NIST. To visualize and analyze the computed networks, the Cytoscape network visualization software (version 3.8.2) (Shannon et al., 2003; Smoot et al., 2011) was used.

All semi-annotated and some unmatched/unidentified nodes, visualized in Cytoscape, were verified using their empirical formulae calculated from accurate mass and fragmentation patterns. In addition to the manual inspection of annotations, some natural products dereplication databases such PubChem (https://pubchem.ncbi.nlm.nih.gov) and Dictionary of Natural Products (http://dnp.chemnetbase.com/faces/chemical/ChemicalSearch.xhtml) were also used. The annotations were also verified and confirmed against available literature. All metabolite annotations were carried out at level 2 and 3 of the Metabolomics Standards Initiative (MSI) (Sumner et al., 2007). All validated annotations (manual and computational) are listed in supplementary table S1. The networks generated were further explored using network annotation propagation (NAP), where first *in silico* fragmentation was applied to the generated FBMN where structural searches were performed in databases *e.g*., GNPS, CHEBI, SUPNAT, and DRUGBANK.

To perform network annotation propagation (NAP) (da Silva et al., 2018) jobs, both the fusion and consensus scores were used based on the first 10 structural candidates. Using scoring methods, NAP re-ranks the candidate structure list based on the network topology (Ernst et al., 2019; Kang et al., 2019; Nephali et al., 2022). The two scoring methods, as previously mentioned, utilized by NAP include (a) fusion scoring, which uses MetFrag *in silico* prediction with MetFusion based on spectral library matches within a molecular family; and (b) consensus scoring, which uses the structural similarity from *in silico* candidates across the spectral nodes of a molecular family. The FBMNs were also explored using DEREPLICATOR (Mohimani et al., 2018) for peptidic structural annotations.

Substructure annotation and initial exploration was carried out using the MS2LDA (MS2 latent Dirichlet allocation) tool (van der Hooft et al., 2016; Rogers et al., 2019) in GNPS and substructure annotations from MotifDB were included in the analyses. From the default settings, some of the parameters were changed as follows for the ESI negative dataset: Bin width was set at 0.01 (Tof data) and LDA free motifs set at 150. The MotifDB included was Rhamnaceae Plant and the rest were excluded. For the ESI positive dataset, the the parameters were changed as follows: LDA free motifs set at 200. The MotifDB included were GNPS, Massbank and Euphorbia. MolNetEnhancer (Ernst et al., 2019) which incorporated outputs from both FBMN and *in silico* tools such as MS2LDA, DEREPLICATOR, NAP and the automated chemical classification through ClassyFire, was applied, providing thus a holistic chemical overview of measured metabolomics spectral data, with enhanced structural details for each fragmentation spectrum. The GNPS job links are provided in the Supplementary materials.

### 2.6 Pathways analysis and relative quantification

The metabolomic pathway analysis (MetPA) tool in MetaboAnalyst, version 5 (Pang et al., 2021) was used to perform functional analysis,

**There are no sources in the current document** particularly pathway analysis. The latter was computed using identifiers of the annotated metabolites, KEGG IDs, as input data. For overrepresentation analysis, the enrichment method, hypergeometric test was used; and for the node importance measure, i.e., topological analysis, the relative betweenness-centrality was employed. For visualization was done using scatterplots and the pathway library used was *Arabidopsis thaliana* (thale cress) (KEGG). Using integrated peak areas, (relative) quantitative analysis was done, and colour-coded heatmap generated in MetaboAnalyst. Pareto-scaling and log-transformation were applied as data pre-treatment methods.

### 2.7 Cell culture

The MIA PaCa-2 pancreatic cancer and HEK 293 human embryonic kidney were cultured in Dulbecco’s Modified Eagle Medium (DMEM) (Sigma Aldrich, USA) supplemented with 10% fetal bovine serum (FBS) (Biogen, UK) and 1% Pen-Strap (penicillin-streptomycin) (Biowhitakker, Germany) at 37 °C in 95% humidity and 5% CO_2_. About 75% of confluent cells were harvested and rinsed with 6 ml Dulbecco’s Phosphate-buffered Saline (DBPS) (Sigma Aldrich, USA). The cells were then trypsinized with 2 ml of 1 X trypsin and were incubated for 45 s (HEK 293) and 90 s (PaCa-2) at 37 °C in 95% humidity and 5% CO_2._ Four millilitres of fresh DMEM were added to both cell lines respectively to stop trypsinization. Into a new 50 ml centrifuge tube, 6 ml of cell solution was transferred into the centrifuge tube, and the cell solution was centrifuged for 4 min at 2200 rpm (779 × *g*). After centrifugation, the pellet was resuspended in 3 ml of DMEM for cell quantification and further subculturing.

After cell culturing, cell quantification was done using trypan blue and TC20 automated cell counter (Bio-Rad). In a 1.5 ml microcentrifuge tube, 10 µl of cells in DMEM was gently mixed with 10 µl of trypan blue and was followed by aliquoting 10 µl of the mixture into the cell counter slide compatible with the TC20. Cell viability and cell concentration were read and cells that exhibited 95% viability were used for cytotoxicity and caspase activity assays.

### 2.8 Extract preparation

The 80% methanol flower extracts of AMNH and RDC were prepared using the protocol reported in **section 2.2**. The 80% methanol plant extracts were then dried into solid using the rotary evaporator and the two extracts (AMNHF and RDCF) were stored at room temperature until further analysis. The dried extracts were made up to a stock concentration of 100 mg/ml w/v in 100% dimethyl sulphoxide (DMSO) (PanReac AppliChem, Germany) respectively. The extract stock solutions were stored in the dark at 4 °C awaiting further analysis.

### 2.9 Alamar blue cellular viability assay

The HEK 293 and MIA PaCa-2 were plated in 96 well plates (at 5000 cells per well). The cells were treated with AMNHF and RDCF and were incubated for 24 hours with an untreated control, the negative control set as 0.1% DMSO and the positive control set as 100 µM etoposide (Sigma Aldrich, Germany), and a media blank column (plated per column). The AMNHF and RDCF extracts were used to treat the cells in a concentration-based series starting from 100 µg/ml and serially diluting it 2X to 3.125 µg/ml. A volume of 10 µl of the Alamar blue (Thermofisher, USA) reagent was added to each well following the 24 hours treatment and the plate was incubated for 2 hours at 37 °C. The fluorescence was measured in each well using a plate reader and the Gen 5 program 530/25 nm excitation and 590/35 nm emission. The analysis was repeated three times.

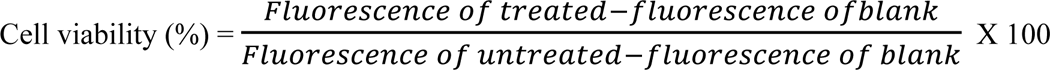

### 2.10 Caspase Glo® 3/7 activity assay

MIA PaCa-2 and HEK 293 cells were plated and treated with AMNHF and RDCF in 33mm petri dishes in a similar manner as described in **section 2.9** with the IC50s and controls. The MIA PaCa-2 cell line was subjected to a caspase detection assay to determine if caspase-3 and −7 were activated due to the compound treatment. The cells were plated for 24 hours in 33mm petri dishes at 1×10^5^ cells/ml. The cells were treated 24 hours after plating with 2 ml treatments consisting of an untreated control, the positive and negative controls, and the AMNHF and RDCF treatments with the IC50s of each extract. The assay was repeated three times and was conducted using the Caspase Glo® 3/7 assay kit (Promega, USA) based on the manufacturer’s recommendations.

### 2.11 In silico molecular docking

The PDB files of the target proteins were obtained from the Protein Data Bank (https://www.rcsb.org/) database and selected based on their resolution: <2.0Å (1.5–2.0Å), while the structures of the ligands were retrieved from PubChem (https://pubchem.ncbi.nlm.nih.gov/). The files were then prepared in UCSF Chimera for docking by deleting and adding hydrogens to the receptors and ligands respectively. The structures were respectively saved in .pdb and .mol2 formats and converted into rec.pdb and .pdbqt formats using Autodock tools. After predicting the active sites of the receptors, a grid box was created to surround the binding residues present in the site using specific X, Y, and Z dimensions and centres. Setting the results to give five prediction outputs, Raccoon and Autodock Graphical user interface supplied by MGL tools were then used to dock the ligands onto the receptors. The complexes exhibiting the lowest Z-scores i.e., the least energy functions were then selected. The resulting .pdb files were viewed in UCSF Chimera, which was used to identify and label the residues involved in the docking.

## 3. Results and Discussion

### 3.1 Chemometric analysis of cannabis cultivars

As detailed in the experimental section, a nontargeted LC-MS/MS-based metabolomics approach was applied for profiling the metabolome of leaves and flowers of two Cannabis cultivars that are used for medicinal purposes: Amnesia haze (*C. sativa* dominant) and Royal dutch cheese (*C. indica* dominant). The methanol extracts from leaf and flower tissues were analyzed on an LC-MS/MS platform. The processed spectral data was explored and evaluated by applying unsupervised chemometrics methods, namely, principal component analysis (PCA) and hierarchical clustering analysis (HCA) modelling. The generated models revealed certain structures within the data, such as tissues-and cultivar-related sample groupings, pointing to underlying differential metabolite profiles which were further investigated (**Figure S1**).

### 3.2. Global chemical profiling – key constituents of cannabis metabolome

The chemical profiling of plant extracts such as Cannabis, achieved through GNPS-based molecular networking (MN) tools, can assist in bettering and laying the foundation for the differentiation of Cannabis cultivars into chemovars based on their chemical class content. In doing so, metabolite biomarkers can be identified for each chemovar, and this would aid in understanding the non-generic medicinal or bio-active potential elicited by each cannabis chemovar or plant-tissue thus narrowing down the medicinal applications of the chemovar of interest. Thus, in this study, the metabolic inventory of the two Cannabis cultivars was investigated. The obtained spectral data were submitted to mass spectral molecular networking (MN) through the GNPS ecosystem, a scalable workflow that digitalizes the diversity and distribution of metabolites in plants (Kang et al., 2019).

Feature-based molecular networking (FBMN) enabled the semi-automated putative annotation of metabolites with confidence level 2 (or level 3) annotations as defined in the proposed minimum reporting standards of the metabolomics standards initiatives (MSI) (Sumner et al., 2007) (**Figure 1**). FBMN represents a computational strategy that facilitates the visualization and compound annotation of complex, high-resolution untargeted LC-MS/MS metabolite data from natural extracts (Hammerle et al., 2021). This type of MN is designed to distinguish structural isomers by incorporating features such as chromatographic retention times which enhances metabolite annotation and thus the dereplication of metabolites whilst also retaining semi-quantitative information to perform statistical analyses (Quinn et al., 2017; Nothias et al., 2018; Nothias et al., 2020). The computed FBMN of spectral data from leaves samples (from both cultivars) comprised 2914 nodes (consensus spectra) where 997 nodes clustered into 128 molecular families based on spectral similarity (**Figure 1.A**). Searching spectral libraries in GNPS, 166 nodes were putatively annotated, with varying relative distributions across the cultivars, which point to diverse chemistries in the leaves of these Cannabis cultivars. For the samples from the flower tissues, the computed FBMN was of 2829 nodes of which 119 nodes were putatively annotated and 998 nodes clustered to form 172 molecular families (**Figure 1.B**). The singletons (selfloop nodes) positioned at the bottom of both networks (**Figures 1.A-B**) represented spectra that were not clustered into molecular families, *i.e*., low spectral similarity scores.

**Figure 1:**
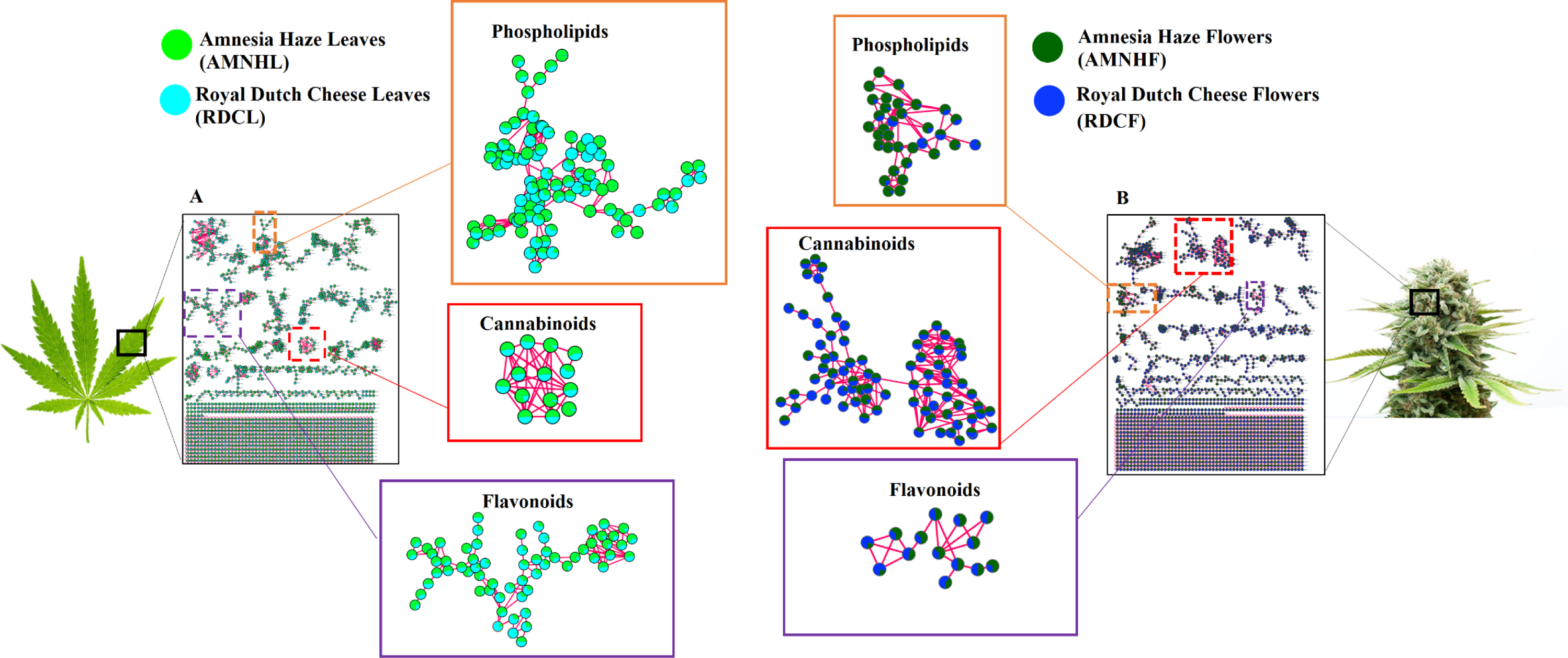
Feature-based molecular networking (FBMN): Molecular networks representing the general mass spectrometry detected chemical space of cannabis cultivars Amnesia haze (AMNH) and Royal dutch cheese (RDC). (A) mass spectral molecular network for the leaves, L denoting leaves and (B) mass spectral network for the flowers, F denoting flowers. **Key**: Amnesia haze leaves (**AMNHL**), Amnesia haze flowers (**AMNHF**), Royal dutch cheese leaves (**RDCL**) and Royal dutch cheese flowers (**RDCF**). Some major molecular families from each network are highlighted, *e.g.,* flavonoids, cannabinoids and phospholipid Clusters.

The molecular families (MFs) in the networks provided a global visualization and insights on the metabolome of the two Cannabis cultivars, revealing diverse metabolite classes including phospholipids, cannabinoids and flavonoids, which are differentially distributed across the cultivars and across the tissues (**Figure 1** and **Figure S2**). From the 128 MFs found in the leaf data (where two or more connected nodes are considered as an MF), 94 MFs were comprised of unknown metabolites. However, 34 MFs had at least one metabolite automatically matched to the GNPS libraries, a similar trend that was also observed in the MFs found in flower MN. Considering that MN is rationally constructed based on spectral similarities (Aron et al., 2020; Vincenti et al., 2020; Yu et al., 2022; Zhang et al., 2023), automated annotation one node in a cluster can aid in decoding and annotating other structurally similar metabolites or features in the same molecular family. As such, MN improves on metabolite annotation, decoding ‘dark matter’ in spectral data, which subsequently provides an improved coverage on the annotated metabolome. In this study, this is illustrated by **Figure 2A** where one spectral node in the MF was putatively annotated through GNPS spectral library matching as CBD. Based on the concept of molecular networking, theoretically this cluster posed as a cannabinoid cluster formed by metabolites with similar structures and fragmentation patterns. This then propagated the manual annotation of unidentified metabolites such as cannabidiolic and cannabidivarinic acids in the cluster (**Figure 2A**). The unannotated metabolites in this cluster also suggest that there may be other unknown CBD related compounds or analogues present.

**Figure 2:**
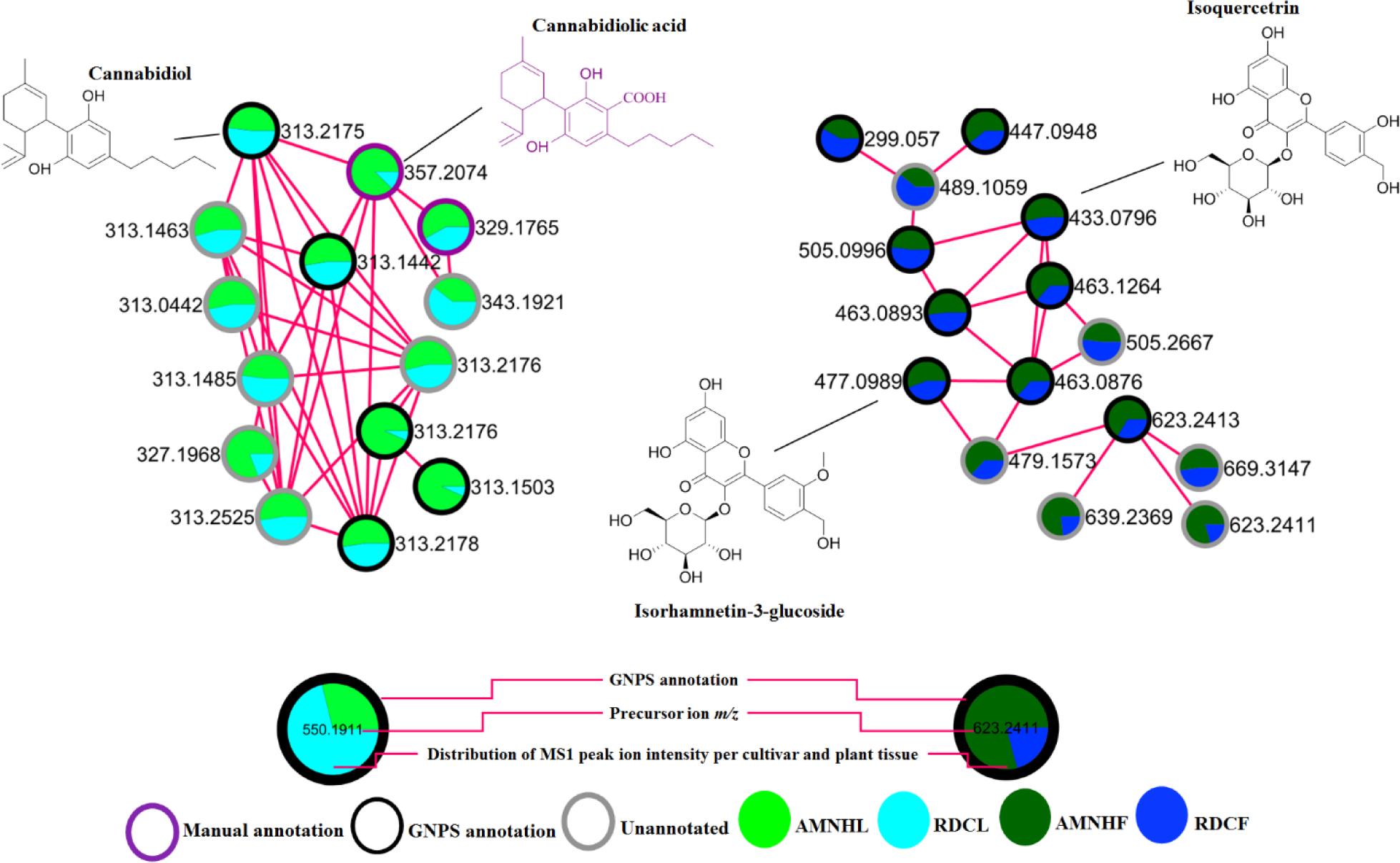
Molecular clusters extracted from FBMN. A detailed visual representation of the output of feature-based molecular networking highlighting some of the major chemical classes in leaves /flowers of Amnesia haze and Royal dutch cheese. **Key**: Amnesia haze leaves (**AMNHL**), Amnesia haze flowers (**AMNHF**), Royal dutch cheese leaves (**RDCL**) and Royal dutch cheese flowers (**RDCF**). (**A**) Cannabinoid cluster in the leaves consisting of cannabidiolic (*m/z* 357.20) and cannabidivarinic acid (*m/z* 329.17) (**B**) flavonoid cluster in the flowers consisting of Isoquecetrin (*m/z* 463.12) and isorhamnetin (*m/z* 477.09). In addition to the annotation of the metabolites in the clusters, FBMN also aids in visualizing the relative abundance of each metabolite in the extracts.

Furthermore, as may be expected from plant samples, flavonoids were also one of the major metabolite classes found in the cannabis tissues. In this flavonoid cluster, there are spectral nodes (such as *m/z* 489.10, 505.28, 479.15, 639.23 and 669.31) that could not be annotated through GNPS spectral library matching, yet they are postulated to be structurally related to glycosylated flavonoids (**Table S1**) due to their grouping with flavonoids such as isoquercitrin (*m/z* 433) and isorhamnetin-3-glucoside (*m/z* 477) (**Figure 2B**)(**Table S1**).

As above mentioned, it is worth noting that applying these computational tools improves annotation as it provides insights into possible molecular families in the measured metabolome. However, there are some limitations, related to similarity scoring algorithms and complexity of untargeted metabolomics spectral data. To improve on this, the “chemical baiting” concept was explored and applied. The baiting approach is a concept that is predominantly used in protein biochemistry where chemical baits or probes are used to bind or concentrate a diverse range of biomarkers (proteins & peptides, metabolites, lipids & fatty acids, nucleic acids, and post translationally modified peptides) that are either occurring in low concentrations or masked by other dominant resident proteins (Luchini et al., 2010). Nonetheless, within metabolomics approaches based on molecular networking (MN), a recognizable exogenous compound (a bait) can be employed to pinpoint other endogenous metabolites residing within the same molecular family cluster. Thus, in this study, the vitexin standard was used as a probe to determine the effect of chemical baiting on FBMN (**Figure 3**).

**Figure 3:**
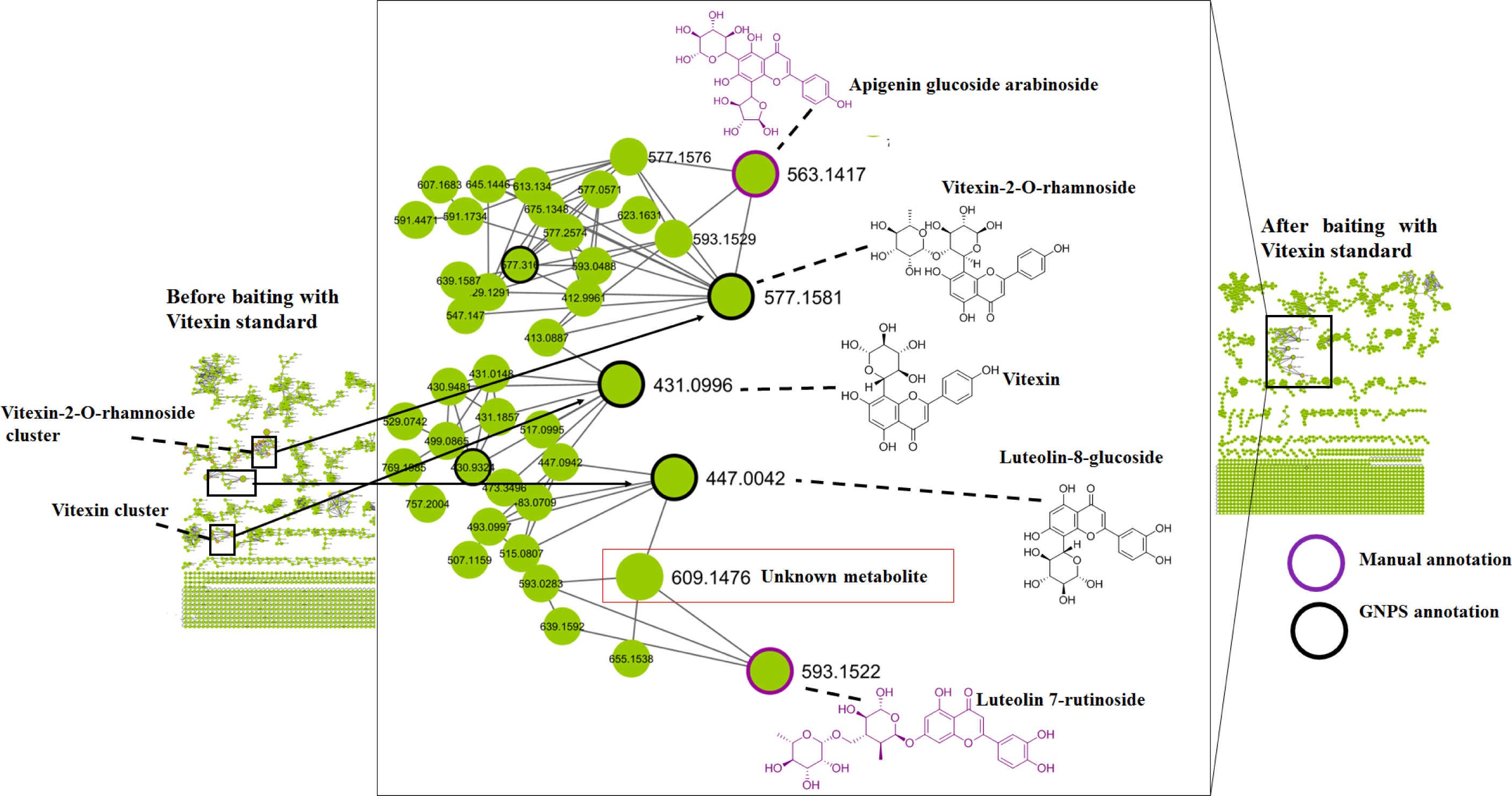
Chemical baiting in FBMN. The effect of chemical baiting on FBMN using the Vitexin standard as a probe for Vitexin-2-O-rhamnoside and any other metabolites that are structurally similar to vitexin. (**A**) FBMN before analyzing samples in the presence of the standard, (**B**) FBMN after analyzing samples with the standard and (**C**) zoomed-in vitexin cluster from (B)-showing the clustering of the vitexin standard together with vitexin-2-O-rhamnoside, as well as the incorporation of an unknown singleton and other structurally similar metabolites into one molecular family.

The overall observed effect of chemical baiting on FBMN (**Figure 3**), showed that using a standard, in this case the vitexin standard, can aid in probing or fishing out known, “unknown knowns” and “unknown unknown” that are structurally similar and thus have similar fragmentation patterns depending on one or more standards used. This was shown by the clustering of vitexin and vitexin-2-O-rhamnoside into one molecular family when the vitexin standard was added as a chemical bait. It is worth noting that cannflavins A and B, flavonoids unique Cannabis cultivars, were also initially detected in the same MF as vitexin-2-O-rhamnoside before the addition of the vitexin standard. However, once the vitexin standard was added, the cannflavins formed their own MF (**Figure 4**). Cannflavins (canflavin A, B and C) are geranylated flavones that are said to be unique to Cannabis cultivars (Rea et al.,2019; Bautista et al., 2021; Tomko et al., 2022).

**Figure 4:**
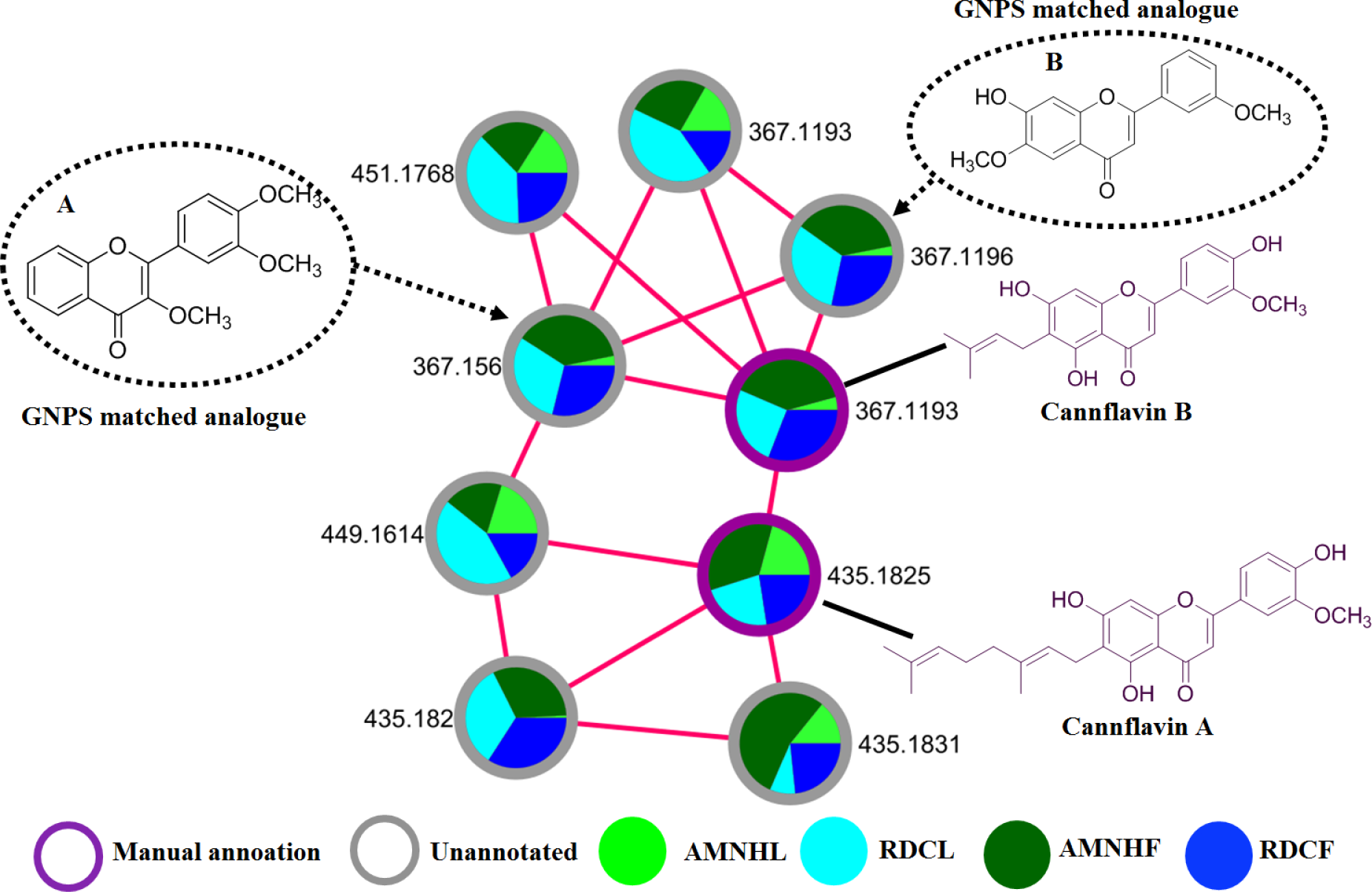
Cannflavin cluster. A representation of the distribution of the detected canflavins (canlfavin A and B) and some of the GNPS matched analogues acquired in ESI negative. **Key**: Amnesia haze leaves (**AMNHL**), Amnesia haze flowers (**AMNHF**), Royal dutch cheese leaves (**RDCL**) and Royal dutch cheese flowers (**RDCF**). (**A**) Uknown compound (*m/z* 367.156) (B) unknown compound (*m/z* 367.1193) matched through GNPS analogue library as afrormosin and ‘3,3’,4’-trimethoxyflavone respectively.

Cannflavins A and B were detected in both the leaves and flowers of AMNH and RDC and reported in **Table S1**. Cannflavin A (*m/z* 435.1825) was distributed evenly across the plant tissues of both cultivars and cannflavin B (*m/z* 367.1193) was recognisably present in the leaves and flowers of RDC but significantly abundant in the flowers of AMNH (**Figure 4**). Using the GNPS analogues library search, the unknown metabolites in the cannflavin cluster were matched to flavone analogs such as afrormosin and ‘3,3’,4’-trimethoxyflavone which are structurally similar to the detected cannflavins. Therefore, this alludes to the notion that combing chemical baiting and molecular networking can help in spectral similarity clustering, and subsequently could increase the confidence in the (semi) automated metabolite annotations. Thus, the computed FBMN after chemical baiting (**Figure 3**) provided spectral clustering and putative annotations of the measured metabolomes from the leaves of the cannabis cultivars, revealing 289 spectral library matches of flavonoids, cannabinoids, phospholipids and more (**Table S1**), 1265 connected nodes and 192 molecular families. Overall, the generated network showed clusters were lipid-like molecules to be more abundant in the plant tissues of RDC and flavonoids such as isoquercitrin to be more abundant in the tissues of AMNH while a cluster of hydroxy acids such as citric acid to occured in similar amounts across both the cultivars (**Figure S2**).

To further explore and characterize the metabolome of the Cannabis cultivars, *in silico* annotation tools such as substructure recognition topic modelling through the MS2 Latent Dirichlet Allocation (MS2LDA), and the network annotation propagation (NAP) were performed for all spectral datasets from both cultivars, AMNH and RDC. To illustrate the contribution of these tools to the Cannabis metabolite annotation and identification at a scaffold diversity level, the outputs obtained from both AMNH and RDC dataset are reported herein (**Figure 5** and **Table S1**). The machine learning (ML)-based tool, MS2LDA, allows an unsupervised decomposition of fragment spectra, discovering patterns of co-occurring fragments and neutral losses (termed ‘Mass2Motifs’, or shortly ‘m2m’) from different MS/MS spectra, which enables extracting information on substructural diversity within each class of metabolites (van der Hooft et al., 2016). Additionally, the generated substructural information highlights the (bio)chemical relationships of the connected compounds. Moreover, these (shared) substructures or scaffolds can point to functional groups and/or core structures of the compounds, revealing common biosynthetic routes in a molecular family (Van Der Hooft et al., 2016; Beniddir et al., 2021; Ramabulana et al., 2021).

**Figure 5.**
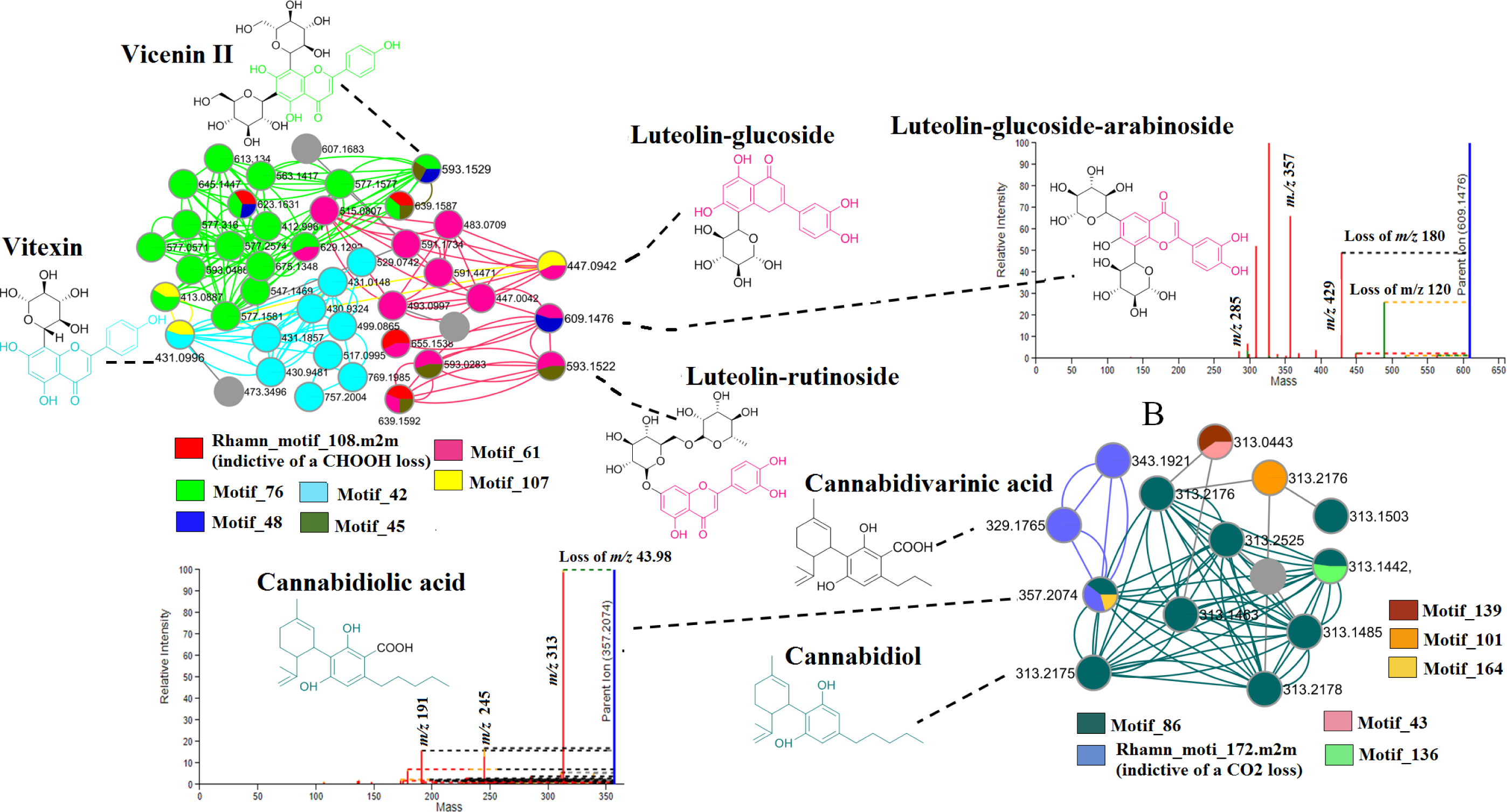
MS2LDA output. molecular network clusters extracted from an MS2LDA substructure exploration analysis of cannabis cultivars AMNH and RDC in ESI negative ionization mode. (**A**) and (**B**) illustrate the MS2LDA-driven annotations of flavonoids (A) and cannabinoids (B) in the leaves of the cultivars. The coloured nodes represent some of the recognized substructures.

One hundred and eighty-one (181) Mass2Motifs were discovered, and most were common in both the Cannabis leaves and flower datasets (ESI negative) of the two cultivars, respectively. The Mass2Motifs, annotated and unannotated were visualized through the MS2LDA website (ms2lda.org) (Wandy et al., 2018). As an illustration, some of the characterized Mass2Motifs found in the Cannabis leaves are highlighted and shown in **Figure 5**. These discovered substructural Mass2Motifs are significant in the sense that they can aid in the structural elucidation of “unknown knowns’’ (metabolites or spectral nodes whose reference spectra are not found in the GNPS spectral libraries and therefore represented as unknowns) such as *m/z* 609.1476 identified as luteolin-glucoside-arabinoside. When evaluating the chemical makeup of the glucosylated flavonoids, **Figure 5.A** illustrates that *m/z* 609.1476 shares a common substructure (unknown motif_61) with the manually annotated metabolites luteolin-glucoside and luteolin-rutinoside. This resulted in the structural elucidation of luteolin-glucoside-arabinoside (*m/z* 609.1476) and led to the shared unknown motif_61 being annotated as a substructure that is related luteolin (*m/z* 285). Additionally, this also suggests that motif_107, motif_48, and motif_45 are all related to some sugar loss, respectively. Moreover, it can be postulated that motif_ 76 and motif_42 are both related to apigenin as they both form the core structure of the putative annotations of vitexin (*m/z* 431) and vicenin II (*m/z* 593) MS2LDA also highlighted some of the chemical moieties that make up cannabinoids as shown in **Figure 5.B**.

Compounds such as cannabidiol (CBD) were annotated through GNPS spectral library matching and with the use of FBMN, it was shown to be structurally similar to cannabidiolic acid (CBDA - *m/z* 357.2074) and cannabidivarinic acid (CBDVA - *m/z* 329.1765) (**Figure 2**). The MS2LDA discovered substructures of these compounds (or cluster) clearly describe the chemical relationship between the compounds through rhamn_motif_172.m2m which describes the carboxylic acid that is attached to CBDA and CBDVA (**Figure 5.B**). Furthermore, this led to the shared unknown motif_86 being annotated as a substructure that is related to cannabidiol (*m/z* 313.2175). Combining Mass2Motif annotations and library matches, we can observe how the known compounds and unkown cannbidiol analogues are associated with the cannabidiol substructure. This makes sense as the cannabidiol derivatives would have a cannabidiol core structure. Altogether, the here highlighted Mass2Motifs are indicative of flavonoids and cannabinoids, respectively, and their decorations (i.e., glycosylations). Consequently, this confirms and gives confidence to the FBMN annotations such as those previously made on the highlighted molecular family (**Figure 2**) and listed in **Table S1.**

In addition to MS2LDA, Network Annotation Propagation (NAP) was also applied. NAP performs *in silico* fragmentation-based metabolite predictions or annotations of candidate structures through searching several databases including GNPS, ChEBI (Chemical Entities of Biological Interest), SUPER NATURAL (II) and DNP. Therefore, using these two scoring methods, NAP is a beneficial tool when experimental spectra are matched to a few spectral libraries matches as it also allows the propagation of annotations in the absence and presence of library spectral matches (da Silva et al., 2018; Nephali et al., 2022; Kim et al., 2023). The The outputs of the NAP obtained in this study are reported in the **supplementary material.** In brief, we could verify that NAP was able to correctly predict 18 (out of 997 spectral features) metabolites for the leaves data and 25 (out of 998) for the flower data. In both leave and flower exctracts, the NAP predicted metabolites include, cannabidiol, vitexin, orientin, and luteolin. These metabolite annotations were verified through manual inspection of fragmentation patterns and literature.

To get an improved chemical compound class annotation and a comprehensive chemical overview of the measured spectral data, the outputs from FBMN, MS2LDA, NAP and DEREPLICATOR+ were combined in an enhanced molecular network workflow, MolNetEnhancer, which integrates also the automated chemical classification of molecular families with ClassyFire (Ernst et al., 2019). MolNetEnhancer enabled the chemical annotation, visualization, and discovery of the subtle substructural diversity within molecular families (**Figure 6**).

**Figure 6:**
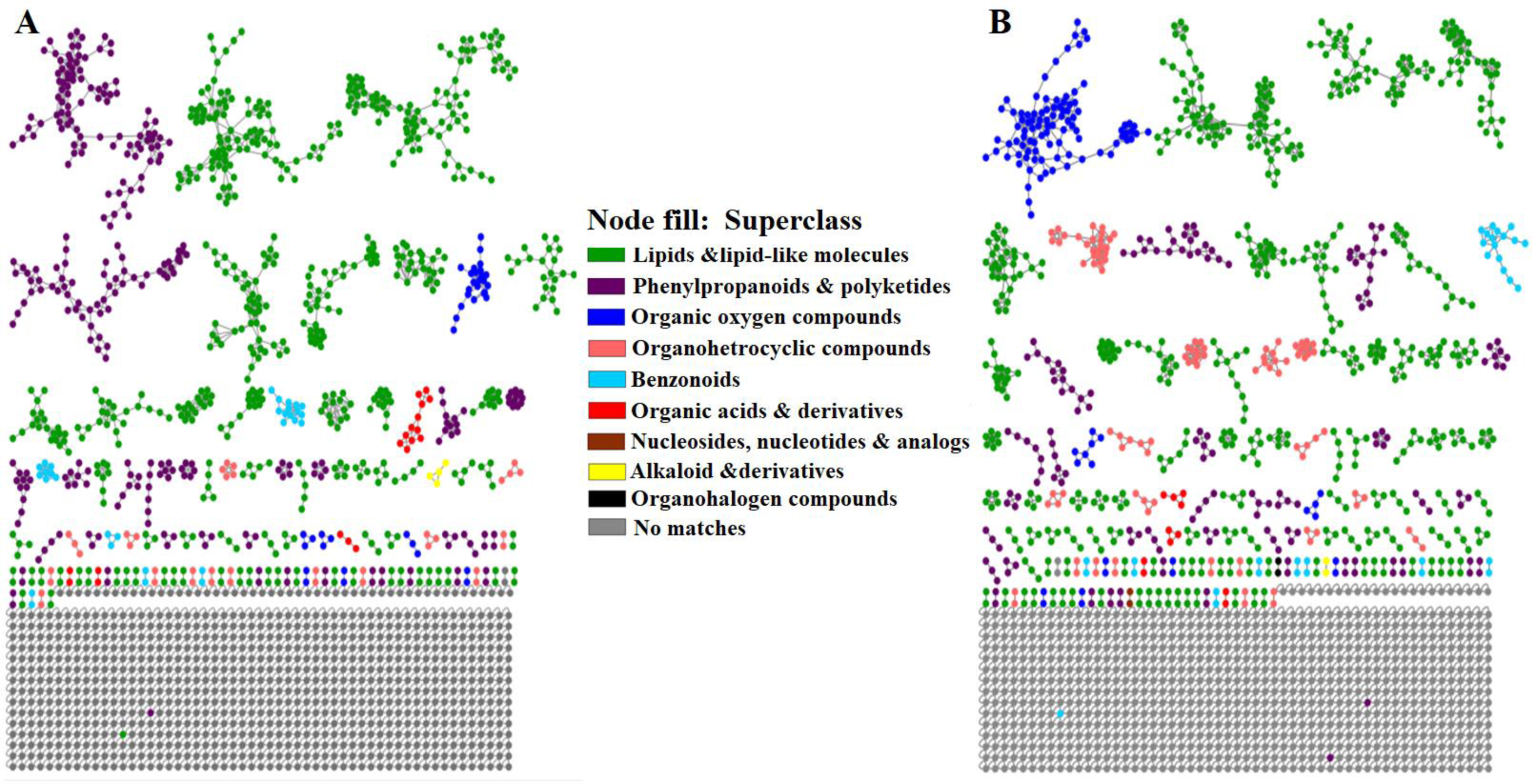
Enhanced molecular networking. MolNetEnhancer mass spectral FBMN of (**A**) leaves and (**B**) flowers of both AMNH and RDC cannabis cultivars acquired in ESI negative. The MolNetEnhancer FBMNs highlight the chemical superclasses present in leaves and flowers of two cultivars. The networks were computed from the combined outputs of FBMN putative annotations, substructure annotations (MS2LDA), network annotation propagation (NAP), and DEREPLICATOR.

The generated networks indicated the varying distribution of major superclasses such as phenylpropanoids & polyketides, lipids and lipid-like molecules, organic oxygen compounds, benzonoids and organoheterocyclic compounds across the leaves (**Figure 6.A**) and flowers (**Figure 6.B)**. A major difference observed between the plant-tissues of the cultivars (**Figure 6**) was the abundance of phenylpropanoids and polyketides superclass, which was the second most dominating superclass in the leaves of the cultivars (**Figure 6.A**). Phenylpropanoids and polyketides include flavonoids, lignins, phenolic acids, stilbenes and coumarins. A considerable number of flavonoids were identified in the leaves and these metabolites, in addition to their plant functions, exhibit biological activities such anti-inflammatory and antioxidant properties.

Other polyphenolic compounds such as feruloyl quinic acid and caffeoylquinic acid reported in **Table S1** were also detected in the leaves. These phenolic acids which form part of chlorogenic acids are known to possess anticarcinogenic properties due to their antioxidant activities (Makhafola et al., 2016; Bouyahya et al., 2022). When evaluating the superclasses present in the flowers of the two cutivars, the Phenylpropanoids and polyketide superclass (comprised of polyphenolic compounds) was less dominant. Contradictory to this observation, flowers are regarded as important reproductive organs thus high levels of polyphenols are expected (Piccolella et al., 2020). However, the clear lack of polyphenols in the flowers of the studied cultivars (**Figure 6.B**) poses questions on the presence of intact biosynthetic routes of these compounds or the effects that the breeding process could have had on the chemistry of these cultivars. Moreover, literature regarding polyphenols in the flowers of Cannabis cultivars remains scarce and limited (Izzo et al., 2020). It has been shown that Cannabis cultivars lose genetic variation due to domestication and excessive breeding for selective traits (Clark & Merlin 2016). Therefore, such losses could contribute to the observed differences in the distribution of the phenylpropanoids and polyketides (polyphenols) between the leaves and flowers of the studied cultivars.

Additionally, Cannabis genomics data lacks plant-tissue-specific data (Hussain et al., 2021) but the metabolite data generated in this study could lay the foundation for further exploration of the regulatory genetic networks of chemical classes in plant tissues of Cannabis cultivars. For example, when zooming in on these identified superclasses and detailing the relative abundance of the identified metabolites, **Figure S2** highlights the cultivar-specific and plant-tissue-specific metabolite differences where lipid-like molecules such as Glc-Glc-octadecatrienoyl-sn-glycerol (isomer 2) are dominant in the tissues of RDC while flavonoids (phenylpropanoids and polyketide superclass) such as isoquercitrin are more dominant in the tissues of AMNH. These metabolic differences could be related to changes in regulatory networks underlying the specialized metabolome of these cultivars.

The chemical profiling achieved through the GNPS-molecular networking tools revealed a complex and diverse phytochemical space of the cultivars based on the leaf and flower metabolite profiles. The leaves and flowers were characterized by the presence of chemical classes such as flavonoids, cannabinoids and phospholipids which have various medicinal properties as discussed above. Possible anti-cancer properties are one of the medicinal properties that were postulated from the revealed metabolomes of the two cultivars. To further explore this postulation, *in silico* molecular docking studies were performed to computationally predict the anti-cancer or anti-proliferative properties of the revealed cultivar chemistries supported by biological assay studies.

### 3.3. Anti-proliferative properties of AMNH and RDC: *In silico* molecular docking and biological assay studies

The phytochemistry of both cultivars illuminated various chemical classes including cannabinoids and flavonoids-with some described to have anti-cancer properties amongst other biological activities. Selected cannabinoid and flavonoid ligands, based on the elucidated chemical profiles of the two cultivars (**Tables S1** and **S3**), were docked against various cancer targets. Each ligand was docked against its known target since the targets have been shown or suggested to play vital roles in cancer pathways. The interactions of the ligands with the receptor/protein targets were investigated through computer-assisted modelling to reveal their effective binding capacities (**Table S3**). Illustratively, the most significant receptor and ligand interactive binding affinity or docking score (−10.3 kcal/mol) was observed between cannabinoid receptor type 2 (CB2) and △^9^*-*THC (**Figure 7)**. The amino acid residues involved in the interaction were shown to be SER285, PHE281, VAL113 and SER90.

**Figure 7:**
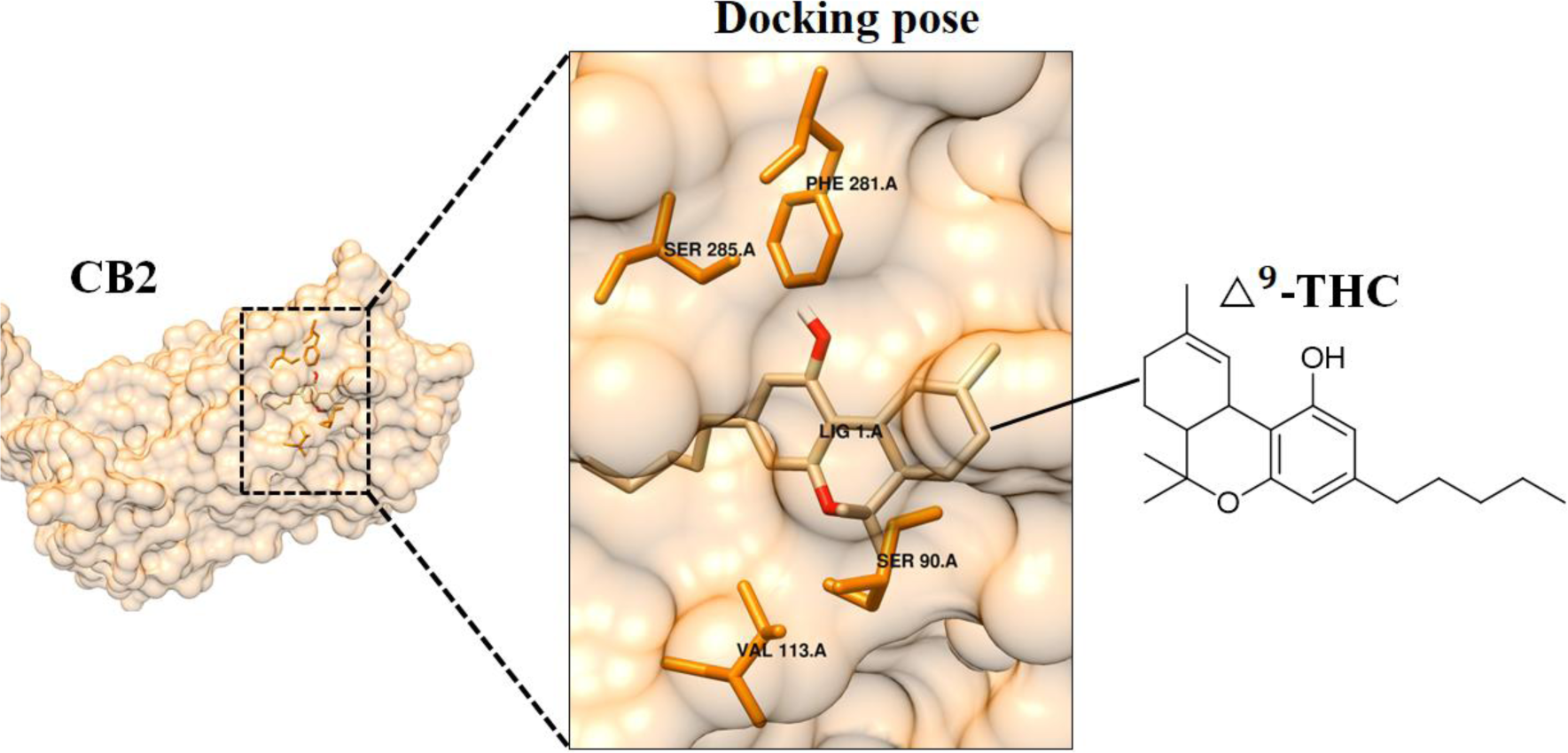
Molecular docking to predict the activity of selected cannabis metabolites. Structure of CB2 active site in complex with △^9^-THC. The docking pose indicates the receptor interactive amino acid residues SER285, PHE281, VAL113 and SER90 that position △^9^-THC in the binding pocket of CB2.

This was followed by the interaction between cannabinoid receptor 2 (CB2) and CBD where the binding affinity of −9.5 kcal/mol.CB1 and CB2 are endocannabinoid cannabinoid receptors or membrane proteins with seven transmembrane helices and belong to the rhodopsin-like G-protein-coupled receptor superfamily (Sharif et al., 2016). CB1 is predominately found in the central nervous system (CNS) and in addition to its ability to bind to a wide range of cannabinoids, CB1 actively binds and mediates the psychoactive effects of △^9^-THC (Kendall & Yudowski, 2017). On the other hand, CB2, which had the highest docking score or interacting with △^9^-THC, is known to play a significant role in the regulation of immune responses, inflammation, pain, and other metabolic processes (Yeliseev & Gawrisch, 2017). In the many types of cancers, the activation of both CB1 and CB2 triggers several pathways. In lung cancer, it has been reported that when bound to the CB receptors, △^9^-THC upregulates Tribbles homolog 3 (TRB3) which propagates autophagy mediated apoptosis (Salazar et al., 2009; Fu et al., 2023). Moreover, studies conducted on pancreatic Mia PaCa2 cell line showed that, △^9^-THC induced caspase-3 activation (characteristic of apoptotic cell death as illustrated above) and stimulated the *de novo* synthesis of ceramide which increased the cell apoptotic rate through the up-regulation of stress-regulated protein p8 (Carracedo et al., 2006; Laezza et al., 2020).

The cannabinoid CBD, which is known to interact with CB1 and CB2 in cancer pathways, was also shown to interact with the G-coupled protein receptor 55 (GPR55) where the docking score was measured to be −7.8 kcal/mol (**Table S3**), slightly lower than the docking scores on CB1 and CB2. GPR55 is one of the newly discovered endogenous cannabinoid receptors that now form part of the extended endocannabinoid system (Ramer et al., 2019). The endocannabinoid receptor GPR55 is abundant in the brain, skeletal muscle, gastrointestinal (GI) tract, white adipose tissue, the islets of Langerhans (β and α cells) and the pancreas. In cancer studies, GPR55 has been investigated for its vital role in cancer-promoting activities (Falasca and Falasca, 2022). The normal stimulation of GPR55 by its endogenous ligand (lysophosphatidylinositol (LPI)) activates the pro-tumorigenic Akt and extracellular receptor kinase (ERK) pathways which promote cell proliferation. However, when CBD binds to GPR55, it acts as an antagonist and thus promotes antiproliferation effects in cancer cell lines (Laezza et al., 2020).

When evaluating the flavonoid content of Cannabis cultivars, literature highlights that more than 20 flavonoids have been identified in Cannabis of which the most abundant are flavone (luteolin and apigenin) and flavonol (quercetin and kaempferol) aglycones and glycosides (Bautista et al., 2021; Cásedas et al., 2022). Thus, flavonoids identified and reported in **Table S1** such as apigenin, quercetin and kaempferol, in their non-glycosylated forms, were also investigated for their binding affinities to cancer targets (**Table S3**). Moreover, the anti-cancer propertie of cannflavin A, a flavone unique to the Cannabis plant, were also investigated. Among all the screened flavonoids, cannflavin A had the most significant docking score of - 9.3 kcal/mol through its interaction with caspase 3 The measured docking score of cannflavin A with caspase 3 amino acids SER144, THR143, GLU196, TRY200, PRO206, TRY202, ARG167 on caspase 3, a cysteine–aspartic acid protease that is predominantly found in the cytoplasm of cells and exists as an inactive pro-enzyme (pro-caspase 3) (Luo et al., 2010; Ponder and Boise, 2019). Studies done on cannflavin A as a ligand to caspase-3 have shown that the interaction causes slight cleavage on caspase-3 which results in cytotoxic effects to a variety of cancer cells such as human bladder transitional carcinoma cells (Tomko et al., 2022). The stimulation of caspase-3 results in apoptosis, is often described as a “point of no return” for a cell and is characterized by apoptotic nuclear changes such as DNA fragmentation, chromatin condensation and nuclear disruption (Luo et al., 2010; Boudreau et al., 2019).

Overall, all the flavonoids and cannabinoids exhibited good binding affinity with the various cancer targets which may be responsible for the anti-proliferative properties of cannabis flower extracts observed in **Figures S4 and Figure 8**. When considering Cannabis in natural products and cancer research, several studies have been done to support the efficacy of Cannabis varieties for various cancer treatment-related symptoms. Some studies have also explored the efficacy of Cannabis in inducing apoptosis in cancer cells. However, most of these studies mainly focus on the effect of isolated major cannabinoids (e.g., CBD, CBG, THC etc) and focus less on other chemical classes or their combined synergistic effect. Since Cannabis flowers are mostly studied in drug discovery and bioactive compound exploration, herein, the flowers of the two Cannabis cultivars (AMNHF and RDCF) previously profiled for their chemical content (through computational metabolomics) were investigated for anti-proliferative effects through *in vitro* cell viability assays complemented with *in silico* molecular docking studies (discussed above) to determine if they elicited similar cytotoxic effects. The *in vitro* cell viability assays such as the alamar blue assay (Longhin et al., 2022) showed that both AMNHF and RDCF extracts were cytotoxic to the cancerous MIA PaCa-2 cell line which indicated anti-proliferative properties (**Figure S4**). The observed anti-proliferative properties were further supported and highlighted by the caspase activities of the MIA PaCa-2 cells upon being treated with AMNHF an RDCF extracts respectively.

**Figure 8:**
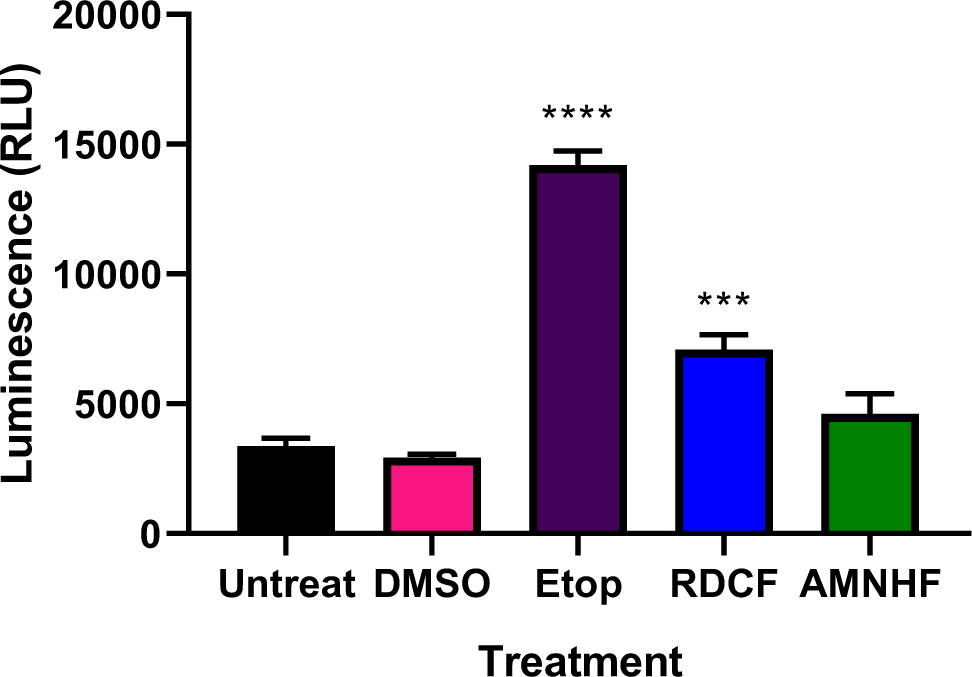
Caspase activity. The average luminescence readings (in relative light units) for the 24-hour treatment of the MIA PaCa 2 cell line showing the untreated cells, the cells treated with 0.1% DMSO, 100 µM etoposide and the IC50s of RDCF (65.7 µg/ml) and AMNHF (2 µg/ml). The asterisk (*) represents (*P < 0.05, **P < 0.01 ***P < 0.001, ****P < 0.0001) as calculated by a T-test between untreated and treated samples.

Various stimuli initiate or induce the energy-dependent apoptosis, an anti-proliferative process, where the activation of enzymes called caspases (cysteine-aspartic proteases) occurs (Saraste & Pulkki, 2000; Elmore, 2007). Caspases are proteolytic enzymes that have well-defined roles in cell death mediated by apoptosis, necroptosis, and autophagy. Apoptotic caspases (involved in both intrinsic and extrinsic apoptotic pathways) are expressed as inactive pro-caspases that are activated to their active forms-caspase 2,3,7,8,9 and 10. Their activation propagates a cascade of signalling events that result in the controlled demolition of cellular components (McIlwain et al., 2013; Shalini et al., 2015).

Cell death caused by AMNHF and RDCF on MIA PaCa-2 treated cells was clearly indicated in **Figure S4**, and since caspases are over-expressed during cell death caused by apoptosis, caspase activity was then investigated. An evident increase in caspase activity was seen for the RDCF and ANMHF treated MIA PaCa-2 cells (**Figure 8**). The untreated cells were observed to have a reading of 3378.3 RLU while RDCF, following the chemotherapeutic drug (etoposide), presented a significantly high average luminescence reading of 7093.3 RLU-which was a 210% increase in caspase activity when compared to the untreated cells. Moreover, the AMNHF treated cells presented an average luminescence reading of 4603.7 RLU, a 136.3% increase in caspase activity which was lower than that of RDCF. However, these evident increases in caspase activity indicate the possible up-regulation of apoptotic caspases (caspase 3 and 7) and thus point to apoptotic events due to the treatment of the cells with RDCF and AMNHF. These results also correlate to the Alamar blue cytotoxic activities observed on the treated MIA PaCa-2 cells.

No data has been reported on the displayed anti-proliferative activities of Cannabis cultivars AMNHF and RDCF. Therefore, we may hypothesize that the observed activities stem from the distinct chemical profiles of the two cultivars. Consequently, AMNHF and RDCF present promising subjects for upcoming investigations, including isolating compounds of various chemical classes and assessing their potential for anti-cancer properties. It is noted that these compounds usually undergo natural metabolism upon introduction into the human body. Therefore, future research focusing on the catabolism of these metabolites following Cannabis consumption and their route within the body will be important to assess the full health-related properties of various Cannabis cultivars.

## 4. Conclusion

The numerous Cannabis cultivars that are made accessible for medicinal purposes, are classified into several categories based only on the plant’s cannabinoid content. Nonetheless, the metabolomics computational strategies used herein such as molecular networking approaches, helped in visualizing, and elucidating the chemical diversity of the studied Cannabis cultivars beyond their cannabinoid content. The revealed plant-tissue based chemical profiles of the studied Cannabis cultivars showed variation in the distribution of phytocannabinoids (major and minor) as well as other phytochemicals. Moreover, the phytochemical space of the leaves (which are underused in clinical studies) was highlighted as an alternative source of compounds with possible medicinal value. Such revelations, including the varying distribution of chemical classes across the studied cultivars, can assist in understanding the metabolome of Cannabis and gives an opportunity for the discovery of novel compounds in the elucidated chemical space.

Since the therapeutic outcomes of Cannabis are non-generic, the knowledge generated from the chemotyping done herein can also inform further experiments such as the testing of the biological activities of the plant-tissues as they may be used to address different ailments based on their non-generic chemical composition. When considering the flower anti-proliferative activities of the extracts, these findings put emphasis on the importance of metabolic profiling Cannabis cultivars as they can inform their use as pharmacological agents, in this case as possible cancer suppressors. However, since this study was based on crude extracts, the anti-proliferative activities observed could be a result of the synergistic effect of multiple compounds. Thus, further studies such as fractionation and compound isolation and evaluation of some of the identified metabolites in the studied Cannabis extracts on cancer cell lines are still needed. Moreover, based on the observed metabolite differences between the plant tissues of the studied cultivars, future studies could include integrating genomics and transcriptomics studies for in-depth analysis of the biosynthetic routes and regulation of the chemical compound classes illuminated in the cultivars.

## Supplementary information

Supplementary data to this article can be found online:

## Acknowledgments

University of Venda, Biochemistry Department are gratefully thanked for access to the SHIMADZU LCMS-9030 qTOF. The South African national research fund (NRF) is highly thanked for bursary support to A.M.

## Author contributions

A.M.: Methodology, data analysis, data curation, writing of original draft, writing-review and editing. M.C.: Supervision, data curation and analysis. A.P.K: Supervision, data curation and analysis. N.E.M.: Supervision, conceptualization, data analysis, data curation, review and editing. J.J.J.vd.H.: Supervision, data curation and analysis, review and editing. F.T: Conceptualization, methodology, data analysis, data curation, writing of original draft, writing-review and editing, funding acquisition and project administration. All authors read and approved the manuscript.

## Data Availability Statement

The spectral data presented in this study is made available on the supplementary files as job links and can also be found in an online repository at https://massive.ucsd. edu/ MSV000093886

