## Supplemental file for "Charting the Cannabis plant chemical space with computational metabolomics"

*Metabolomics*
Supplementary Figures, Tables and GNPS job links

### Charting the Cannabis plant chemical space with computational metabolomics

**Akhona Myoli[1], Mpho Choene[1], Abidemi Paul Kappo[1], Ntakadzeni Edwin Madala[2], Justin J.J. van der Hooft[1,3*], Fidele Tugizimana[1,4,5*]**

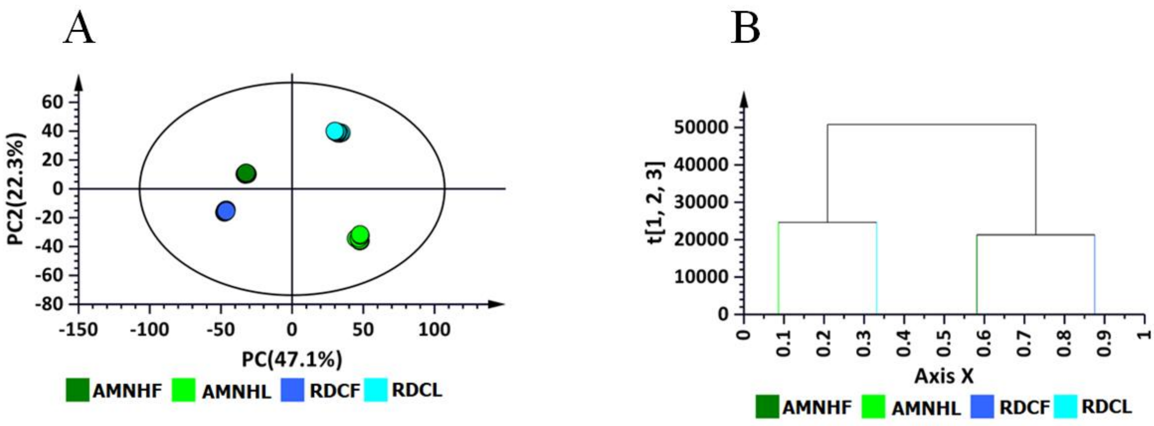


**Figure S1**: **Chemometrics modelling**. Unsupervised analysis and models of the ESI negative data of cannabis cultivars-Amnesia haze and Royal dutch cheese. **Key**: Amnesia haze leaves (**AMNHL**), Amnesia haze flowers (**AMNHF**), Royal dutch cheese leaves (**RDCL**) and Royal dutch cheese flowers (**RDCF**). (**A**) PCA scores plot of the Pareto-scaled dataset of the two cultivars and (**B**) the HCA dendrogram indicating the hierarchical structure of the dataset. The PCA score plot explains R2X = 87.8% variation and Q2 = 86% predictive variation and the HCA corresponds to the variation shown on the PCA scores plot based on seven-fold cross-validation.

#### GNPS molecular networking outputs (job links) for leaves (AMNHL and RDCL) and flowers (AMNHF and RDCF):

FBMN of leaves ESI (-): ID=8cd7fb7a5f5c40e3881566fd2f11bc7d
https://gnps.ucsd.edu/ProteoSAFe/status.jsp?task=8cd7fb7a5f5c40e3881566fd2f11bc7d

FBMN of leaves ESI (+): ID=0e4fabaf54af45a9bc436f209efe5290
https://gnps.ucsd.edu/ProteoSAFe/status.jsp?task=0e4fabaf54af45a9bc436f209efe5290

FBMN of flowers ESI (-): ID=0cd602dc592f460c97a5111d2000a5eb
https://gnps.ucsd.edu/ProteoSAFe/status.jsp?task=0cd602dc592f460c97a5111d2000a5eb

FBMN of flowers ESI (+): ID=6b7f1c180143486692d975e285dcbd58
https://gnps.ucsd.edu/ProteoSAFe/status.jsp?task=6b7f1c180143486692d975e285dcbd58

Combined FBMN ESI (-): ID=a268bab486d5414cb0e716f66fd37781
https://gnps.ucsd.edu/ProteoSAFe/status.jsp?task=a268bab486d5414cb0e716f66fd37781

FBMN of leaves + vitexin standard ESI (-): ID=607fce1557fb46afbeadc69883f5f0df
https://gnps.ucsd.edu/ProteoSAFe/status.jsp?task=607fce1557fb46afbeadc69883f5f0df

FBMN of flowers + vitexin standard ESI (-): ID=9155f3807fd745788840e2315d1192a1
https://gnps.ucsd.edu/ProteoSAFe/status.jsp?task=9155f3807fd745788840e2315d1192a1

DEREPLICATOR + ESI (-) of leaves: ID=943eba0bdc69478b863c22afb7b365dd
https://gnps.ucsd.edu/ProteoSAFe/status.jsp?task=943eba0bdc69478b863c22afb7b365dd

DEREPLICATOR + ESI (+) of leaves: ID=3f00c9f413a9402c9bf713a2b294bd2c
https://gnps.ucsd.edu/ProteoSAFe/status.jsp?task=3f00c9f413a9402c9bf713a2b294bd2c

DEREPLICATOR + ESI (-) of flowers: ID=d6d817f90bc9494e9c5e0f87d477bd1c
https://gnps.ucsd.edu/ProteoSAFe/status.jsp?task=d6d817f90bc9494e9c5e0f87d477bd1c

DEREPLICATOR + ESI (+) of flowers: ID=9ad720b8ec3c40d5abda2615c98c3365
https://gnps.ucsd.edu/ProteoSAFe/status.jsp?task=9ad720b8ec3c40d5abda2615c98c3365

NAP ESI (-) of leaves: ID=378071d9403f44418b7594d6a91d4237
https://gnps.ucsd.edu/ProteoSAFe/status.jsp?task=378071d9403f44418b7594d6a91d4237

NAP ESI (+) of leaves: ID=28f6d06b4cf84950971645ab4fbf281f
https://gnps.ucsd.edu/ProteoSAFe/status.jsp?task=28f6d06b4cf84950971645ab4fbf281f

NAP ESI (-) of flowers: ID=4e851eb2d2524496b1a65c492eeeb784
https://gnps.ucsd.edu/ProteoSAFe/status.jsp?task=4e851eb2d2524496b1a65c492eeeb784

NAP ESI (+) of flowers: ID=7011cdd48fb74bb487ce4db82e8bcb3a
https://gnps.ucsd.edu/ProteoSAFe/status.jsp?task=7011cdd48fb74bb487ce4db82e8bcb3a

MS2LDA ESI (-) of leaves: ID=3ccc5c471c3041c1acc3fce8fc397856
https://gnps.ucsd.edu/ProteoSAFe/status.jsp?task=3ccc5c471c3041c1acc3fce8fc397856

MS2LDA ESI (+) of leaves: D=8cc3b956bcd24c15bcd00f0be4afc17c
https://gnps.ucsd.edu/ProteoSAFe/status.jsp?task=8cc3b956bcd24c15bcd00f0be4afc17c

MS2LDA ESI (-) of flowers: ID=1684184ee24f47af9b812f12addaf821
https://gnps.ucsd.edu/ProteoSAFe/status.jsp?task=1684184ee24f47af9b812f12addaf821

MS2LDA ESI (+) of flowers: ID=f71509557b4942b1ab147ed843fb63d7
https://gnps.ucsd.edu/ProteoSAFe/status.jsp?task=f71509557b4942b1ab147ed843fb63d7

MS2LDA ESI (-) of leaves + vitexin standard: ID=29027ad86fa84025aa73df572862beae
https://gnps.ucsd.edu/ProteoSAFe/status.jsp?task=29027ad86fa84025aa73df572862beae

MS2LDA ESI (-) of flowers + vitexin standard: ID=52bc3bbf53504151acfab5e167a3ef60
https://gnps.ucsd.edu/ProteoSAFe/status.jsp?task=52bc3bbf53504151acfab5e167a3ef60

MolNetEnhancer ESI (-) of leave: ID=d65b3a5b5e7949b28f3ac672b59b2c60
https://gnps.ucsd.edu/ProteoSAFe/status.jsp?task=d65b3a5b5e7949b28f3ac672b59b2c60

MolNetEnhancer ESI (-) of flowers: ID=e9634b2290f04e1bbd976a2e99866822
https://gnps.ucsd.edu/ProteoSAFe/status.jsp?task=e9634b2290f04e1bbd976a2e99866822

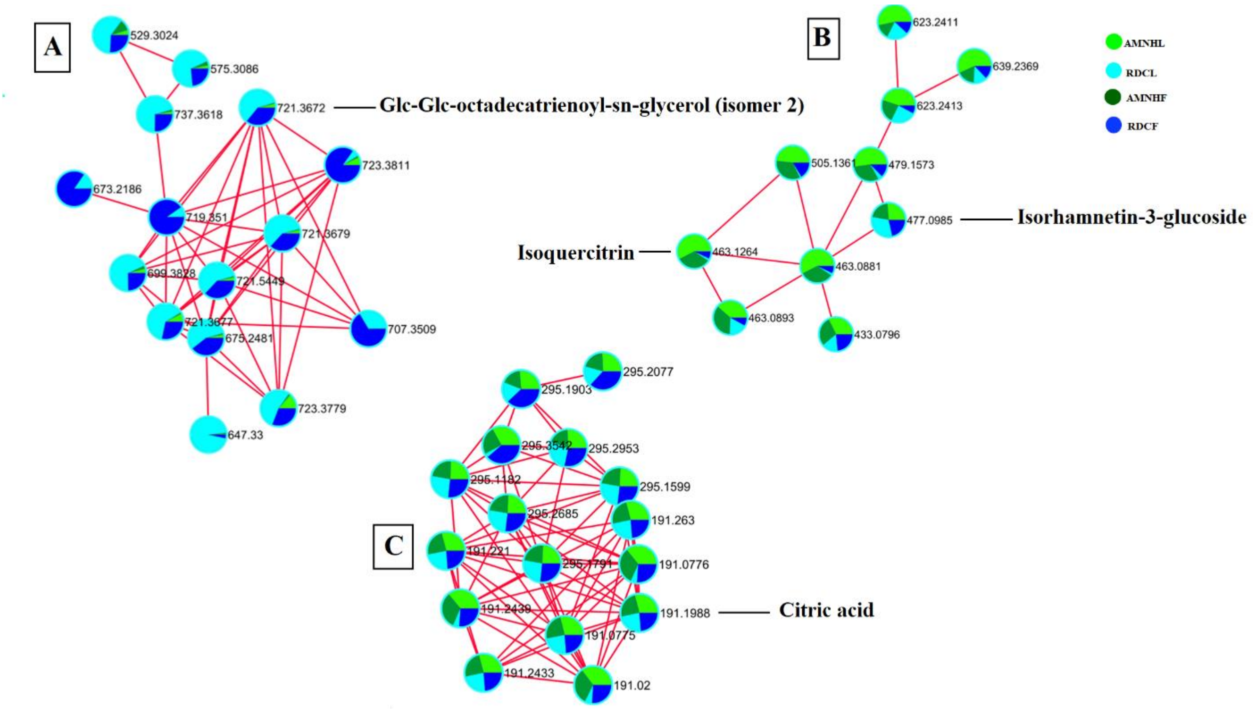


**Figure S2**: **Cannabis cultivar-specific and plant-tissue-specific metabolite differences**. A combined molecular network showing the relative quantification of some metabolites identified across the cultivars and their plant tissues. **Key**: Amnesia haze leaves (**AMNHL**), Amnesia haze flowers (**AMNHF**), Royal dutch cheese leaves (**RDCL**) and Royal dutch cheese flowers (**RDCF**). (A) Shows lipid-like molecules to be more abundant in the plant tissues of RDC, (B) shows flavonoids such as isoquercitrin to be more abundant in the tissues of AMNH while (C) shows a cluster of hydroxy acids such as citric acid to occur in similar amounts across both the cultivars.

**Table S1**: All putatively annotated metabolites (level 2 and 3 MSI) in the plant-tissues of Amnesia haze (AMNH) and Royal dutch cheese (RDC) Presence (✓) and absence (×) stating the distribution of metabolites and (*) denotes the abbreviations used in **Figure S2**.

| No. | *m/z* | Rt | Adduct | Chemical formular | Fragment ions | Putative annotations | *Abbrev or alternative name | Superclass | Amnesia haze (AMNH) | | Royal dutch cheese (RDC) | |
|---|---|---|---|---|---|---|---|---|---|---|---|---|
| | | | | | | | | | Leaves | Flowers | Leaves | Flowers |
| **1** | 166.0854 | 2.422 | $[M+H]^+$ | $C_9H_{11}NO_2$ | 120,107,103 | Phenylalanine | L-Phe | Organic acids and derivatives | ✓ | ✓ | ✓ | ✓ |
| **2** | 205.0961 | 4.562 | $[M+H]^+$ | $C_{11}H_{12}N_2O_2$ | 170,159,143, 132,118 | L-Tryptophan | L-Trp | Organic acids and derivatives | ✓ | ✓ | ✓ | ✓ |
| **3** | 313.218 | 13.01 | $[M-H]^-$ | $C_{21}H_{30}O_2$ | 311,245,191, 136 | Cannabidiol | CBD | lipid and lipid-like molecules | ✓ | ✓ | ✓ | ✓ |
| **4** | 389.198 | 10.48 | $[M-H]^-$ | $C_{22}H_{32}O_6$ | 313,331,229, 205 | Cannabitriolic acid | CBTA | lipid and lipid-like molecules | ✓ | ✓ | ✓ | ✓ |
| **5** | 285.041 | 7.0955 | $[M-H]^-$ | $C_{22}H_{32}O_6$ | 175,151,133 | Luteolin | Luteolin | Phenylpropanoids &polyketides | ✓ | ✓ | ✓ | ✓ |
| **6** | 431.1 | 6.897 | $[M-H]^-$, $[M+H]^+$ | $C_{21}H_{20}O_{10}$ | 117,283,161 | vitexin | Vit | Phenylpropanoids &polyketides | ✓ | ✓ | ✓ | ✓ |
| **7** | 329.183 | 11.66 | $[M-H]^-$ | $C_{20}H_{26}O_4$ | 311,217,199, 107 | Cannabidi-varinic acid | CBDVA | lipid and lipid-like molecules | ✓ | ✓ | ✓ | ✓ |
| **8** | 357.208 | 16.50 | $[M-H]^-$ | $C_{22}H_{30}O_4$ | 311,245,191,1 36 | Cannabidiolic acid | CBDA | lipid and lipid-like molecules | ✓ | ✓ | ✓ | ✓ |
| **9** | 375.2188 | 11.412 | $[M-H]^-$ | $C_{22}H_{30}O_5$ | 357,273,179,1 35,122 | 6,7-epoxy-Cannabigerolic acid | 6,7-Epoxy-CBGA | lipid and lipid-like molecules | ✓ | ✓ | ✓ | ✓ |

| | | | | | | | | | | | | |
|---|---|---|---|---|---|---|---|---|---|---|---|---|
| **10** | 315.0732 | 3.6545 | $[M-H]^{-}$ | $C_7H_6O_4$ | 152,109,108 | 2,5-Dihydroxybenzoic acid | 2,5-Dihydro-acid | Organic acids and derivatives | ✓ | ✗ | ✓ | ✗ |
| **11** | 721.3702 | 11.874 | $[M-H]^{-}$ | $C_{33}H_{56}O_{14}$ | 675,415,397, 277,235 | Glc-Glc-octadecatrienoyl-sn-glycerol (isomer 2) | Glc-Glc-glycerol | lipid and lipid-like molecules | ✓ | ✓ | ✓ | ✓ |
| **12** | 371.187 | 14.01 | $[M-H]^{-}$ | $C_{23}H_{32}O_4$ | 353,327,259, 191 | Cannabidiolic acid monomethyl ether | CBDMA | lipid and lipid-like molecules | ✓ | ✓ | ✓ | ✓ |
| **13** | 459.0943 | 8.783 | $[M-H]^{-}$ | $C_{22}H_{20}O_{11}$ | 113,283,268, 175 | Oroxindin | Oroxindin | Phenylpropanoids &polyketides | ✓ | ✗ | ✓ | ✗ |
| **14** | 312.13 | 7.6065 | $[M-H]^{-}$ | $C_{18}H_{19}NO_2$ | 148,176,190 | Feruloyl-tyramine | Ferlulo-tyr | Phenylpropanoids &polyketides | ✓ | ✓ | ✓ | ✓ |
| **15** | 447.09 | 6.547 | $[M-H]^{-}$ | $C_{21}H_{20}O_{11}$ | 445 | Orientin | Orientin | Phenylpropanoids &polyketides | ✓ | ✓ | ✓ | ✓ |
| **16** | 299.0566 | 7.8436 | $[M-H]^{-}$ | $C_{16}H_{12}O_6$ | 285,284 | Diosmetin | Diosmetin | Phenylpropanoids &polyketides | ✓ | ✗ | ✓ | ✗ |
| **17** | 577.1533 | 6.9476 | $[M-H]^{-}$ | $C_{27}H_{30}O_{14}$ | 413,293,173 | Vitexin-2-O-rhamnoside | Vit-2-O-rhamn | Phenylpropanoids &polyketides | ✓ | ✓ | ✓ | ✓ |
| **18** | 593.9265 | 6.329 | $[M-H]^{-}$ | $C_{27}H_{30}O_{15}$ | 503,473,383, 353 | Vicenin 2 | Vic-2 | Phenylpropanoids &polyketides | ✓ | ✗ | ✓ | ✗ |
| **19** | 452.2905 | 12.601 | $[M+H]^{+}$ | $C_{21}H_{44}NO_7P$ | 313 | 1-Palmitoyl-2-hydroxy-sn-glycero-3-phosphoethanolamine | 1-Pal-hydr-gly-phos | lipid and lipid-like molecules | ✓ | ✗ | ✓ | ✗ |
| **20** | 597.3066 | 12.99 | $[M-H]^{-}$ | $C_{27}H_{51}O_{12}P$ | 315,281,241,154 | 1-(9Z-Octadecenoyl)-sn-glycero-3-phospho-(1'-myo-inositol) | 9Z-Octa-1-myo-inositol | lipid and lipid-like molecules | ✓ | ✗ | ✓ | ✗ |

| | | | | | | | | | | | | |
|---|---|---|---|---|---|---|---|---|---|---|---|---|
| **21** | 509.2904 | 13.126 | $[M\text{-}H]^-$ | $C_{24}H_{47}O_9P$ | 281,227,153 | 1-(9Z-Octadecenoyl)-sn-glycero-3-phospho-(1'-sn-glycerol) | 9Z-Octa-1-sn-glycerol | lipid and lipid-like molecules | ✓ | ✗ | ✓ | ✗ |
| **22** | 571.2912 | 12.864 | $[M\text{-}H]^-$ | $C_{25}H_{49}O_{12}P$ | 391,315,255, 241,153 | 1-Hexadecanoyl-sn-glycero-3-phospho-(1'-myo-inositol) | 1-Hexa-phopho-1-myo-inositol | lipid and lipid-like molecules | ✓ | ✗ | ✓ | ✗ |
| **23** | 493.1 | 6.611 | $[M+FA\text{-}H]^-$ | $C_{21}H_{20}O_{11}$ | 447,357,327 | Luteolin-6-c-glucoside | Lut-6-glu | Phenylpropanoids &polyketides | ✓ | ✓ | ✓ | ✓ |
| **24** | 483.2749 | 12.698 | $[M\text{-}H]^-$ | $C_{22}H_{45}O_9P$ | 255,245,227, 153 | 1-Hexadecanoyl-sn-glycero-3-phospho-(1'-sn-glycerol) | 1-Hexa-phopho-1-sn-gly | lipid and lipid-like molecules | ✓ | ✓ | ✓ | ✓ |
| **25** | 437.1823 | 9.908 | $[M+H]^+$, $[M\text{-}H]^-$ | $C_{26}H_{28}O_6$ | 313,289,299 | Cannflavin A | Cannflavin A | Phenylpropanoids &polyketides | ✓ | ✓ | ✓ | ✓ |
| **26** | 549.2557 | 7.307 | $[M+FA\text{-}H]^-$ | $C_{24}H_{40}O_{11}$ | 527 | 2-Cyclohexen-1-one, 3,5,5-trimethyl-4-[3-[(6-O-beta-D-xylopyranosyl-beta-D-glucopyranosyl)oxy]butyl] | 2-cycl-1-one | lipid and lipid-like molecules | ✓ | ✓ | ✓ | ✓ |
| **27** | 353.0889 | 5.113 | $[M\text{-}H]^-$ | $C_{16}H_{18}O_9$ | 179,191,135 | Caffeoylquinic acid | Caffeo-qui-acid | Phenylpropanoids &polyketides | ✓ | ✓ | ✓ | ✓ |
| **28** | 591.1811 | 7.4177 | $[M\text{-}H]^-$ | $C_{28}H_{32}O_{14}$ | 504,427,367, 326,307 | 2-O-rhamnosyl-swertisin | 2-O-rhamn-swert | Phenylpropanoids &polyketides | ✓ | ✗ | ✓ | ✗ |

| | | | | | | | | | | | | |
|---|---|---|---|---|---|---|---|---|---|---|---|---|
| **29** | 327.2178 | 9.546 | $[M-H]^-$ | $C_{18}H_{32}O_5$ | 211,229,183, 171 | 9,12,13-trihydroxyoctadeca-10,15-dienoic acid | Trihydroxy-octa-dienic-acid | lipid and lipid-like molecules | ✓ | ✓ | ✓ | ✓ |
| **30** | 367.1447 | 7.1555 | $[M-H]^-$ | $C_{17}H_{20}O_9$ | 191,173,193, 134 | Feruloyl quinic acid (isomer of 887, 888) | Ferul-qui-acid | Phenylpropanoids &polyketides | ✓ | ✓ | ✓ | ✓ |
| **31** | 433.0768 | 7.234 | $[M-H]^-$ | $C_{20}H_{18}O_{11}$ | 300,271,255,301 | Guaijaverin | Guaijaverin | Phenylpropanoids &polyketides | ✓ | ✓ | ✓ | ✓ |
| **32** | 609.146 | 7.01 | $[M-H]^-$ | $C_{27}H_{30}O_{16}$ | 300,271,151 | Quercetin-3-rutinoside | Quercetin-3-rut | Phenylpropanoids &polyketides | ✓ | ✓ | ✓ | ✓ |
| **33** | 709.3739 | 13.118 | $[2M-2H+Na]^-$ | $C_{18}H_{16}O_7$ | 343,299,301 | Usnic acid | Usnic acid | Benzonoids | ✗ | ✓ | ✗ | ✓ |
| **34** | 773.3542 | 11.315 | $[2M-Na]^-$ | $C_{19}H_{20}O_8$ | 375 | Pseudo placodiolic acid | Pseudo-plac-acid | Benzonoids | ✗ | ✓ | ✗ | ✓ |
| **35** | 577.1346 | 5.557 | $[M-H]^-$ | $C_{30}H_{26}O_{12}$ | 451,425,407,289,161 | Procyanidin B1 | Procyanidin B | Phenylpropanoids &polyketides | ✓ | ✓ | ✓ | ✓ |
| **36** | 452.28 | 12.268 | $[M-H]^-$ | $C_{21}H_{44}NO_7P$ | 255,213 | Phosphatidylethanolamine | Lyso 16:0 | lipid and lipid-like molecules | ✓ | ✓ | ✓ | ✓ |
| **37** | 477.1033 | 7.677 | $[M-H]^-$ | $C_{22}H_{22}O_{12}$ | 315,314,285 ,271 | Isorhamnetin-3-glucoside | Isorhamn-3-glu | Phenylpropanoids &polyketides | ✓ | ✓ | ✓ | ✓ |
| **38** | 289.07 | 5.976 | $[M-H]^-$ | $C_{15}H_{14}O_6$ | 287,245,137,125,109 | (-)-Epicatechin | Epicatechin | Phenylpropanoids &polyketides | ✓ | ✓ | ✓ | ✓ |
| **39** | 463.09 | 7.219 | $[M-H]^-$ | $C_{21}H_{20}O_{12}$ | 301,271,255 | Isoquercitrin | Isoquercitrin | Phenylpropanoids &polyketides | ✓ | ✓ | ✓ | ✓ |
| **40** | 579.1498 | 5.997 | $2[M-H]^-$ | $C_{15}H_{14}O_6$ | 289.,245,125 | Catechin dimer | 2XCatechin | Phenylpropanoids &polyketides | ✓ | ✓ | ✓ | ✓ |

| | | | | | | | | | | | | |
|---|---|---|---|---|---|---|---|---|---|---|---|---|
| **41** | 223.097 | 9.534 | $[M-H]^-$ | $C_{12}H_{16}O_4$ | 179,137 | Olivetolic acid | Olivet-acid | Phenylpropanoids &polyketides | × | ✓ | × | ✓ |
| **42** | 553.1658 | 9.634 | $[M-H]^-$ | $C_{25}H_{30}O_{14}$ | 391,373,347 | Ligustrosidic acid | Ligus-acid | lipid and lipid-like molecules | × | ✓ | × | ✓ |
| **43** | 403.1309 | 6.819 | $[M-H]^-$ | $C_{17}H_{24}O_{11}$ | 101,113,119,179,223 | Oleoside 11-methyl ester | Oleo-methy-ester | lipid and lipid-like molecules | × | ✓ | × | ✓ |
| **44** | 617.1451 | 6.685 | $[M-Na]^+$ | $C_{27}H_{30}O_{15}$ | 455,437,599 | Saponarin | Saponarin | Phenylpropanoids &polyketides | ✓ | ✓ | ✓ | ✓ |
| **45** | 601.1513 | 6.847 | $[M-Na]^+$ | $C_{30}H_{26}O_{12}$ | 583,481,455 | Vitexin 2"-O-p-coumarate | Vit-2-O-coumarate | Phenylpropanoids &polyketides | ✓ | ✓ | ✓ | ✓ |
| **46** | 524.3703 | 13.216 | $[M+H]^+$ | $C_{26}H_{54}NO_7P$ | 524,184,104 | 1-Octadecanoyl-sn-glycero-3-phosphocholine | 1-Octadec-gly-phospho | lipid and lipid-like molecules | ✓ | ✓ | ✓ | ✓ |
| **47** | 461.074 | 7.139 | $[M-H]^-$ | $C_{21}H_{18}O_{12}$ | 285,229 | Kaempferol 3-glucuronoside | Kaemp-3-glu | Phenylpropanoids &polyketides | ✓ | ✓ | ✓ | ✓ |
| **48** | 518.3212 | 12.699 | $[M+Na]^+$ | $C_{24}H_{50}NO_7P$ | 459,313,184,146,104 | 1-Hexadecanoyl-sn-glycero-3-phosphocholine | 1-Hexadec-gly-phospho | lipid and lipid-like molecules | ✓ | ✓ | ✓ | ✓ |
| **49** | 311.2076 | 13.98 | $[M+H]^+$ | $C_{21}H_{26}O_2$ | 223,121,109 | Cannabinol | CBN | lipid and lipid-like molecules | ✓ | ✓ | ✓ | ✓ |
| **50** | 341.2101 | 17.01 | $[M+H]^+$ | $C_{22}H_{29}O_3$ | 341,285,219,161 | Tetrahydrocannabinolic acid-acylium ion ($\triangle^9$-THCA-1) | $\triangle^9$-THCA-1 | lipid and lipid-like molecules | ✓ | ✓ | ✓ | ✓ |
| **51** | 341.2104 | 12.04 | $[M+H]^+$ | $C_{22}H_{29}O_3$ | 229,219,161,117 | Cannabidiolic acid- actrylium ion (CBDA-1) | CBDA-1 | lipid and lipid-like molecules | ✓ | ✓ | ✓ | ✓ |
| **52** | 359.2209 | 17.38 | $[M+H]^+$ | $C_{22}H_{30}O_4$ | 285,261,219,135 | Canna-bicycliolic acid | CBLA | lipid and lipid-like molecules | ✓ | ✓ | ✓ | ✓ |
| **53** | 463.3242 | 7.593 | $[M+H]^+$ | $C_{21}H_{18}O_{12}$ | 343,313,287 | Breviscapine | Breviscapine | Phenylpropanoids &polyketides | ✓ | ✓ | ✓ | ✓ |
| **54** | 338.412 | 14.336 | $[M+H]^+$ | $C_{22}H_{43}NO$ | 149,163,135,121,114, 107 | Erucamide | Erucamide | lipid and lipid-like molecules | ✓ | × | ✓ | × |

| | | | | | | | | | | | | |
|---|---|---|---|---|---|---|---|---|---|---|---|---|
| **55** | 353.2079 | 12.151 | $[M+H]^+$ | $C_{21}H_{36}O_4$ | 261,243,173,149,121,107 | Monolinolenin | Monolin | lipid and lipid-like molecules | ✓ | ✓ | ✓ | ✓ |
| **56** | 315.231 | 15.43 | $[M+H]^+$ | $C_{21}H_{30}O_2$ | 193,135,123,109,107 | $\triangle^9$−Tetrahydro cannabinol | $\triangle^9$-THC | lipid and lipid-like molecules | ✕ | ✓ | ✕ | ✓ |
| **57** | 355.1897 | 15.93 | $[M+H]^+$ | $C_{22}H_{26}O_4$ | 337,313,253,219 | Cannabinolic acid | CBDA | lipid and lipid-like molecules | ✓ | ✓ | ✓ | ✓ |
| **58** | 522.3542 | 12.708 | $[M+H]^+$ | $C_{26}H_{52}NO_7P$ | 504,184,125,104 | 1-oleoyl-glycero-3-phosphochloline | 1-Oleo-gly-3-phospho chloline | lipid and lipid-like molecules | ✕ | ✓ | ✕ | ✓ |
| **59** | 601.13 | 5.271 | $[M+Na]^+$ | $C_{30}H_{26}O_{12}$ | 311,449,313 | Procyanidin B2 | Procy-B2 | Phenylpropanoids &polyketides | ✕ | ✓ | ✕ | ✓ |
| **60** | 796.45 | 14.938 | $[M+H]^+$ | $C_{42}H_{66}O_{14}$ | 735,517 | Tenacissoside H | Tena-H | lipid and lipid-like molecules | ✓ | ✓ | ✓ | ✓ |
| **61** | 395.26 | 11.614 | $[M+H]^+$ | $C_{19}H_{22}O_9$ | 377,359,352 | Aloesin | Aloesin | Organoherocyclic compounds | ✕ | ✓ | ✕ | ✓ |
| **62** | 413.2298 | 11.772 | $[M+Na]^+$ | $C_{23}H_{34}O_5$ | 395,377,231,149 | Gitoxigenin | Gitox | lipid and lipid-like molecules | ✓ | ✓ | ✓ | ✓ |
| **63** | 219.1733 | 9 | $[M+Na]^+$ | $C_{12}H_{20}O_2$ | 159,145,133,131,119,105 | Linalyl butyrate | Lin-buty | Organic acids and derivatives | ✕ | ✓ | ✕ | ✓ |
| **64** | 439.2295 | 7.331 | $[M+H]^+$ | $C_{21}H_{26}O_{10}$ | 277,259,203 | Monnieriside G | Monn-G | Phenylpropanoids &polyketides | ✓ | ✓ | ✓ | ✓ |
| **65** | 563.5502 | 13.1255 | $2[M+H]^+$ | $C_{18}H_{35}NO$ | 282,265,247,135,109 | 9-Octadecenamide | 9-Octade | lipid and lipid-like molecules | ✕ | ✓ | ✕ | ✓ |
| **66** | 182.0805 | 1.276 | $[M+H]^+$ | $C_9H_{11}NO_3$ | 147,136,123,119,107, 103 | L-Tyrosine | L-Tyr | Organic acids and derivatives | ✓ | ✓ | ✓ | ✓ |
| **67** | 369.110 | 11.712 | $[M+H]^+$ $[M-H]^-$ | $C_{21}H_{20}O_6$ | 313,298,165 | Cannflavin B | Cannflavin-B | Phenylpropanoids &polyketides | ✓ | ✓ | ✓ | ✓ |

| | | | | | | | | | | | | |
|---|---|---|---|---|---|---|---|---|---|---|---|---|
| **68** | 563.1410 | 5.855 | $[M-H]^-$ | $C_{26}H_{28}O_{14}$ | 449,281,223, 163,135 | Apigenin glucoside arabinoside | Api-glu-arab | Phenylpropanoids &polyketides | ✓ | ✓ | ✓ | ✓ |
| **69** | 593.1520 | 6.553 | $[M-H]^-$ | $C_{27}H_{30}O_{15}$ | 473,429,327,309 | Luteolin-7-rutinoside | Lut-7-rut | Phenylpropanoids &polyketides | ✓ | ✓ | ✓ | ✓ |
| **70** | 609.1470 | 6.651 | $[M-H]^-$ | $C_{27}H_{30}O_{16}$ | 489,429,327, 309 | Luteolin-glucoside-arabinoside | Lut-glu-arab | Phenylpropanoids &polyketides | ✓ | ✓ | ✓ | ✓ |
| **71** | 299.057 | 9.09 | $[M-H]^-$ | $C_{16}H_{12}O_6$ | 285,284,256 | Chrysoeriol | Chrysoeriol | Phenylpropanoids &polyketides | ✗ | ✓ | ✗ | ✓ |
| **72** | 191.1980 | 1.02 | $[M-H]^-$ | $C_6H_8O_7$ | 147, 129, 87 | Citric acid | Citric acid | Organic acids and derivatives | ✓ | ✓ | ✓ | ✓ |

**Table S2**: Significant metabolic pathways in cannabis cultivars Amnesia haze (AMNH) and Royal dutch cheese (RDC) based on the KEGG IDs of metabolites identified in the leaves and flowers of the cultivars.

| No. | Pathway | Total | Hits | *p*-value | Impact |
|---|---|---|---|---|---|
| **1** | Flavone and flavonol biosynthesis | 10 | 2 | 8.4457E-4 | 0 |
| **2** | Flavonoid biosynthesis | 47 | 3 | 9.5604E-4 | 0.02046 |
| **3** | Phenylalanine, tyrosine, and tryptophan biosynthesis | 22 | 2 | 0.0042195 | 0.02002 |
| **4** | Phenylpropanoid biosynthesis | 46 | 2 | 0.017904 | 0.03161 |
| **5** | Stilbenoid, diarylheptanoid and gingerol biosynthesis | 8 | 1 | 0.03723 | 0.13235 |
| **6** | Tyrosine metabolism | 16 | 1 | 0.073265 | 0.10811 |
| **7** | Tryptophan metabolism | 28 | 1 | 0.12515 | 0.12307 |
| **8** | Isoquinoline alkaloid biosynthesis | 6 | 1 | 0.028036 | 0.5 |
| **9** | Aminoacyl-tRNA biosynthesis | 46 | 2 | 0.017904 | 0 |
| **10** | Indole alkaloid biosynthesis | 4 | 1 | 0.018766 | 0 |
| **11** | Glycine, serine, and threonine metabolism | 33 | 1 | 0.14602 | 0 |
| **12** | Ubiquinone and other terpenoid-quinone biosynthesis | 38 | 1 | 0.16646 | 0 |
| **13** | Glucosinolate biosynthesis | 65 | 1 | 0.26976 | 0 |

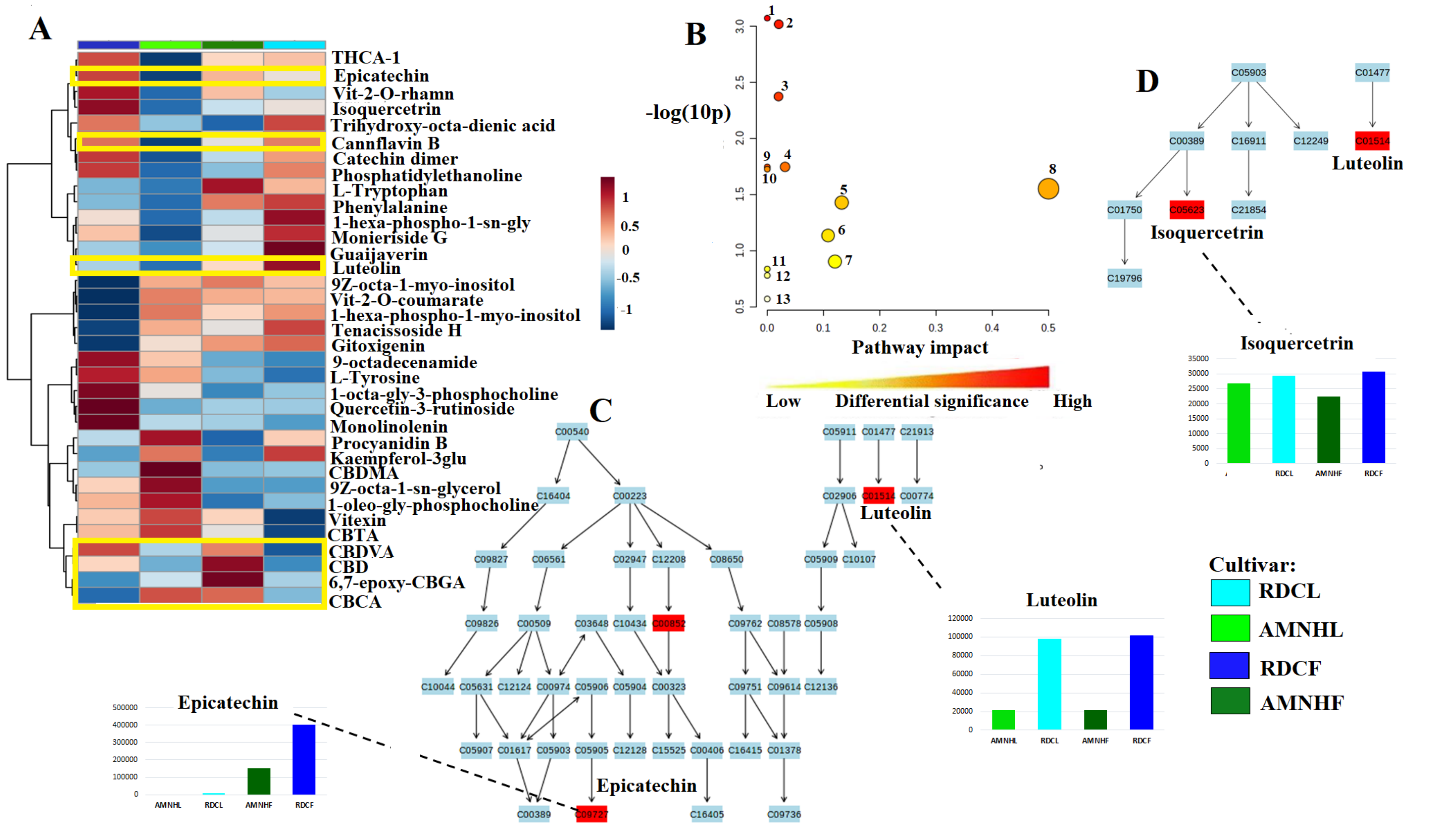


**Figure S3**: **Functional analysis**. Relative quantification of some metabolites identified across the leaves and flowers of AMNH and RDC cultivars (A). Summary of the metabolism pathways identified in the cannabis cultivars (B) where (C) represents the flavonoid and (D) flavone and flavanol biosynthesis pathways.

**Table S3**: Binding interactions identified for some cannabinoids and flavonoids ranked according to the docking score (binding affinity energy (kcal/mol), where a low docking score indicates good/significant interaction and a high score indicating bad/insignificant interaction between protein target and ligand).

| Target and Ligand | Interacting Residues | Docking Score |
|---|---|---|
| **Cannabinoid receptor 2 (CB2) & △$^9$−tetrahydrocannabinol (△$^9$-THC)** | SER285 ,PHE281, VAL113, SER90 | **-10.3 kcal/mol** |
| **Cannabinoid receptor 1 (CB1) & Cannabidiol (CBD)** | PHE108, PHE268, MET384, MET103, PHE170. | **-9.5 kcal/mol** |
| **Caspase 3 & Cannflavin A** | SER144, THR143, GLU196, TRY200, PRO206, TRY202, ARG167 | **-9.3 kcal/mol** |
| **Cannabinoid receptor 1 (CB1) & △$^9$−tetrahydrocannabinol (△$^9$-THC)** | PHE108, PHE268, SER383, MET103, PHE170 | **-9.2 kcal/mol** |
| **Cannabinoid receptor 2 (CB1) & Cannabidiol (CBD)** | HIS95, PHE91, PHE87, LYS109, PHE106, ILE110 | **-9.2 kcal/mol** |
| **Mitogen-Activated Protein Kinase 1 (MEK1) & Apigenin** | TRP275, LEU294, LEU244, ILE243, ILE240, GLN236, PHE288, TRP212 | **-8.3 kcal/mol** |
| **G-coupled receptor 55 (GPR55) & Cannabidiol** | VAL103, LEU96, LEU72, LEU100 | **-7.8 kcal/mol** |
| **Fas receptor & Quercetin** | GLU270, ASN268, GLU163, LYS152, ASN154, SER153 | **-6.6 kcal/mol** |
| **Vascular endothelial growth factor (VEGF) & Kaempferol** | LYS286, ASP276, ILE46, ILE48, SER50, ASP34, PHE36 | **-6.6 kcal/mol** |
| **Tyrosine kinase & Luteolin** | ARG184, LYS182, TYR181, TRY209, GLY215, ILE193, SER194, ARG196 | **-6.0 kcal/mol** |

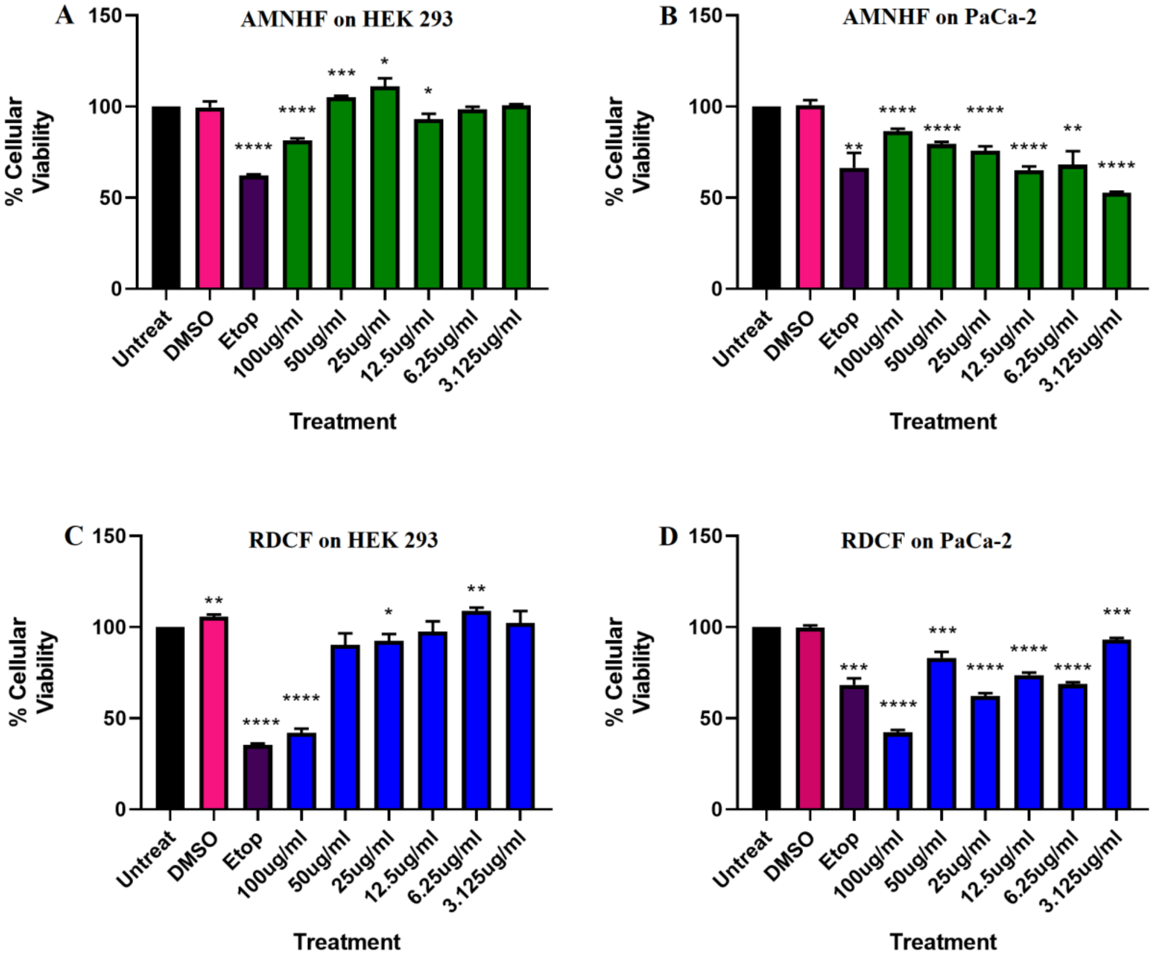


**Figure S4**: **Cell proliferation assessed by Alamar analysis**: (**A**) Percentage viability of HEK 293 (781.7 µg/ml) and (**B**) PaCa-2 cells (IC50 of 3 µg/ml) following the 24-hour treatment with AMNHF respectively. (C) Percentage viability of HEK 293 (IC50 92.48 µg/ml and (D) PaCa-2 cells (IC50 of 65.17 µg/ml) following the 24-hour treatment with RDCF respectively A T-test was performed to assess for statistical significance compared to the positive control (etoposide, p* < 0.05).
